# Sex-linked genomic variation reveals reproductive skew and male-biased dispersal in polygynous sea lions

**DOI:** 10.64898/2026.11.06.764841

**Authors:** Aliya Yakupova, Camille Kessler, Lara Baggio, Stefanie Grosser, Claire R. Peart, Fidel Botero-Castro, Amy B. Baird, John W. Bickham, Joseph I. Hoffman, Jochen B. W. Wolf

## Abstract

Sex-biased dispersal and the resulting asymmetry in gene flow are important factors in evolution. However, their quantification in natural populations remains challenging, not least because sex-specific reproductive skew, as reflected in the mating system, further influences the relative contributions of males and females to the genetic pool of subsequent generations. Incorporating information from mitochondria (MT) and sex chromosomes (X, Y) alongside the more commonly used autosomal markers (AUT) provides a promising framework to partly disentangle confounding processes. Here, we exemplify this approach in two closely related, polygynous species with observational evidence for male-biased dispersal, the Galápagos sea lion (GSL) and the Steller sea lion (SSL). Using whole-genome resequencing data, we leveraged genetic variation among all four genomic compartments, representing genetic variation with female-only (MT), mixed (X, AUT) and male-only (Y) inheritance, to obtain information on the mating system and test for sex-specific migration across geographic and environmental boundaries. For GSL, genomic signatures were consistent with mild polygyny and a moderately elevated rate of male migration, particularly across habitat boundaries. In SSL, male reproductive skew was substantially greater, and male-biased migration was more pronounced and less habitat-dependent. Our results align with the species’ life history and support a link between mating system and dispersal, but were also impacted by confounding biological processes and bioinformatic choices. This study highlights the value and challenges of population genomic approaches integrating multiple genomic compartments to infer variation in individual behaviours.

## Introduction

In animals, sex-biased dispersal translates sex-specific behavioural decisions into the genomic and spatial distribution of genetic variation (de Vries et al., 2025; Sielezniew et al., 2025), with important consequences for adaptation (Platt et al., 2026), speciation (Fraïsse & Sachdeva, 2020; Merondun et al., 2025), and conservation (Brand et al., 2026). When dispersing individuals successfully reproduce, they exchange alleles between populations in proportion to the migration rate *m*. Sex-bias in dispersal can cause migration rates to differ between females and males and introduce an asymmetry in population connectivity among ‘genomic compartments’ differing in their ploidy, mode of transmission, mutational properties and expected effective population sizes (Catalán et al., 2025; Dahms & Momigliano, 2026; Gao et al., 2025; R. J. Petit & Excoffier, 2009). In mammalian systems with XY sex determination, uniparentally inherited components have a direct relationship to migration patterns of each sex: maternally inherited mitochondrial DNA (MT) reflects female-mediated gene flow (*m^F^*), whereas the paternally inherited Y chromosome (Y) reflects male-mediated gene flow (*m^M^*). Autosomes (AUT) are transmitted equally by both sexes and therefore reflect sex-averaged gene flow ((*m^F^ + m^M^*)/2), whereas the X chromosome (X) puts a weight on female-mediated gene flow, as successful female migrants always contribute an X chromosome to the receiving population, whereas males only contribute an X chromosome in half of the cases (*m^F^ + ½ m^M^*). Asymmetry in estimates of gene flow among genomic compartments can thus, in principle, provide a long-term record of sex-biased dispersal complementing traditional inferences based on tracking data collected over just a few years (Petit et al., 2002; Prugnolle & de Meeus, 2002; Shafer et al., 2016). For instance, under male-biased dispersal, which is characteristic of many mammals (Greenwood, 1980), differences in inheritance among genomic compartments are expected to generate the following gradient of population connectivity based on sex-specific differences in migration (commonly quantified by *F*-statistics such as *F_ST_*): strongest differentiation in MT reflecting female philopatry, elevated in X reflecting reduced paternal contribution, intermediate in sex-averaged AUT and weakest in Y reflecting male-mediated gene flow ((MT > X > AUT > Y; **Figure 1A**). However, this pattern is partly confounded by differences in the effective size (*Ne*) among genomic compartments. Because under standard Wright-Fisher assumptions, expected *Ne* is smaller in mitochondria and the Y chromosome (∼1/4 *Ne^AUT^*), intermediate in the X chromosome (∼3/4 *Ne^AUT^*) and largest for autosomes (1 *Ne^AUT^*), genetic drift is expected to be strongest in the former two compartments. Consequently, genetic differentiation is predicted to follow a similar gradient even in the absence of sex-bias in dispersal (MT > Y > X > AUT) making comparisons between mitochondrial and Y-chromosomal differentiation particularly informative for detecting sex-biased dispersal.

**Figure 1.**
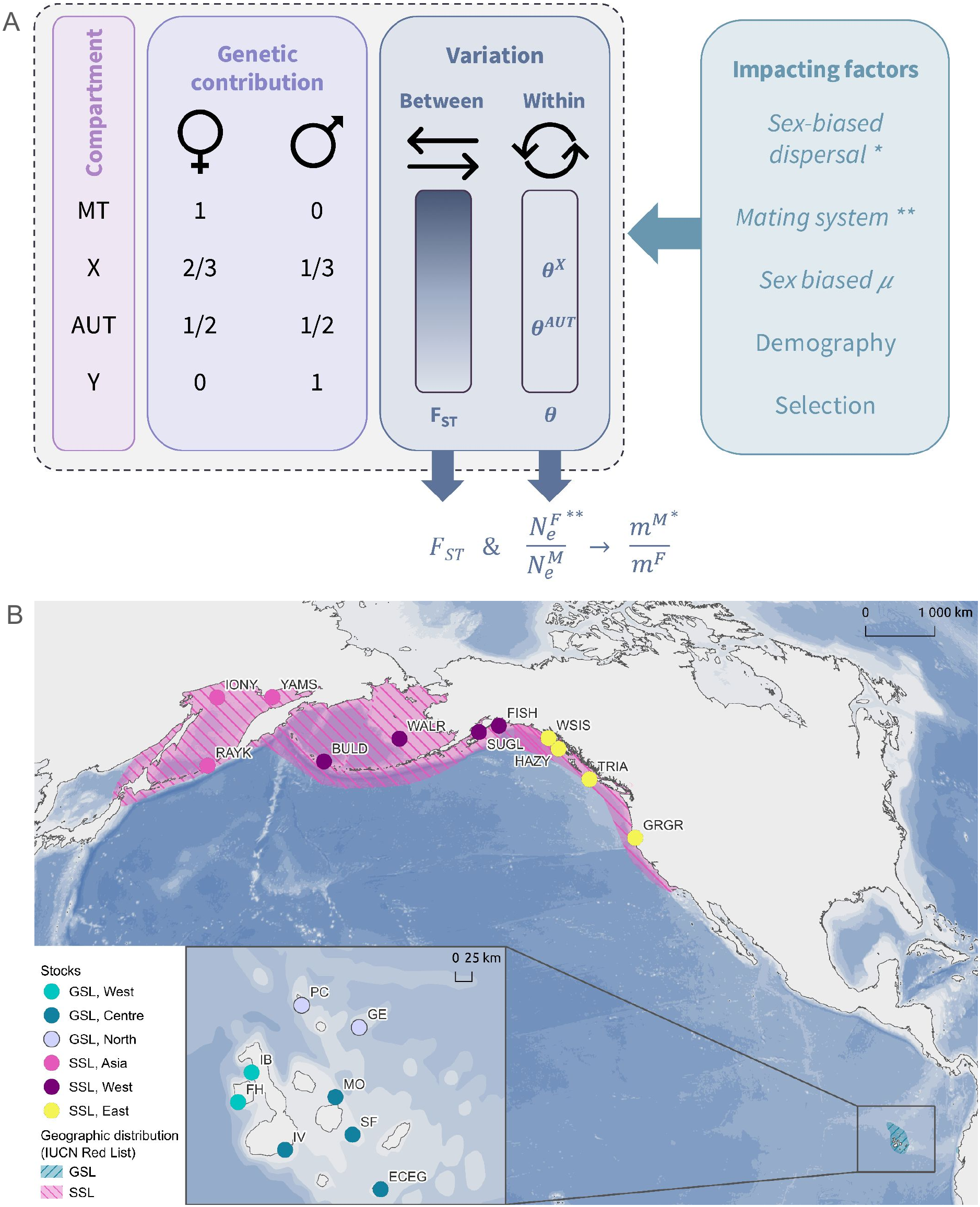
Experimental design. A) Schematic illustrating the inference of sex-bias in reproductive skew and migration based on theoretical expectations for genetic variation across genomic compartments (MT, X, AUT, Y). *From left to right:* The expected contribution of each sex to genetic variation. Genetic differentiation between populations or stocks (*F_ST_*) and genetic diversity within populations (θ) across different compartments; expected relative magnitudes of *F_ST_* are indicated by the gradient, with darker colours representing higher expected values. The rightmost panel lists the most prominent factors impacting genetic variation across genomic compartments. Italicised factors are explicitly taken into account in the calculations of sex-biased reproductive skew and migration. In this study, we use estimates of *F_ST_* and θ across genomic compartments to estimate the ratio of male to female effective population sizes (*Ne^F^/Ne^M^*) reflecting the mating system, and the ratio of male to female migrant rates (*m^M^/m^F^*) to gain insight into sex-bias in dispersal. B) Geographic distribution from the IUCN Red List of GSL (blue) and SSL (pink). Sampling locations are indicated by circles coloured by stock with labels corresponding to rookery names (see **Table S1** for abbreviations).

Sex-biased dispersal, however, often covaries with other life-history traits that can confound patterns of genetic variation, making the interpretation all but straightforward (de Oliveira et al., 2021; Lawson Handley & Perrin, 2007; Li & Kokko, 2019). In mammals, for instance, male-biased dispersal tends to be associated with polygyny (Dobson, 1982; Li & Kokko, 2019). In polygynous species, the effective population size is expected to be lower in males (*Ne^M^*) than in females (*Ne^F^*) because polygyny produces greater variance in male reproductive success (Nunney, 1993). Such mating system induced differences in effective population size among sexes, in turn, alter the number of effective migrants (*m^M^*, *m^F^*) across genomic compartments (**Figure 1A**). Interpretation is further complicated by several other processes that can differentially shape genetic variation across genomic compartments. An increased rate of mutation in the male germline (male-biased mutation rate α) elevates genetic diversity in genomic regions that are inherited through males (Y > AUT > X > MT) (Wilson Sayres, 2018; **Figure S1**), and demographic perturbations can alter the distribution of allelic variation in a compartment-specific manner (Pool & Nielsen, 2007; **Figure S2**). Linked selection can further modify patterns of differentiation, depending on the interaction between the distribution of fitness effects, recombination rate and dominance of mutations differing between haploid MT and Y, hemizygous X and diploid AUT (Botero-Castro & Wolf, 2026; Burri, 2017; Charlesworth, 2012). Moreover, these processes operate over different evolutionary timescales and may therefore contribute to genomic patterns to varying degrees depending on the rate at which mating systems and sex-biased dispersal evolve. Given the complexity of these interacting processes, disentangling the genomic signatures of sex-biased dispersal from the mating system and other confounding factors has proved challenging (Hammer et al., 2008; Laso-Jadart et al., 2025; Spurley & Payseur, 2025).

Here, we seek to assess the degree of sex-biased dispersal while at least partially accounting for the confounding factors mentioned above. To aid this inference, we explore the potential of including the mitochondria and the Y chromosome, which provide a particularly informative contrast between male-and female-mediated gene flow. Historically, the male-limited Y chromosome (or W in ZW systems) has received comparatively less attention in population genetics due to challenges in genome assembly that have only recently been overcome (de Jong et al., 2025; Laso-Jadart et al., 2025; Merondun et al., 2024; Petit et al., 2002; Rhie et al., 2023). Moreover, its highly repetitive sequence content (Gao et al., 2025; Hedrick et al., 2013), duplicated genes and gametologs can compromise reliable inference of genetic variation due to mapping and genotyping artifacts (Greminger et al., 2010; Merondun et al., 2024; Tomaszkiewicz et al., 2016).

As study species, we chose two closely related, highly mobile marine mammals from the Northern Pacific Ocean: (**Figure 1B**): Galápagos sea lions (*Zalophus wollebaeki*; GSL) and Steller sea lions (*Eumetopias jubatus*; SSL). They represent classic examples of species with polygynous mating systems and presumed male-biased dispersal, where adult males roam among rookeries in search of mating opportunities, while females remain philopatric to their natal rookeries (Campbell et al., 2008; Fabiani et al., 2003; Hastings et al., 2017; Lopes et al., 2015; Raum-Suryan et al., 2002). Long-term dispersal data are, however, limited, and dispersal estimates are mostly based on observational data or isotope analysis (Aurioles-Gamboa et al., 2009; Hastings et al., 2017; Wolf et al., 2007; Wolf & Trillmich, 2007), with GPS data limited to short-term foraging behavioural studies (Blakeway et al., 2021; Jeglinski et al., 2015; Lander et al., 2020; McHuron et al., 2016).

Despite the GSL’s ability to disperse across the Galápagos archipelago within a matter of days, prior studies have identified two distinct genetic clusters, with the strongest differentiation between colonies of the western and eastern islands (Jeglinski et al., 2015; Wolf et al., 2008). In SSL, previous studies identified three stocks, with the Eastern stock consistently ranked as most divergent, while more gene flow appears to persist between the Asian and Western stocks (Baker et al., 2005; Harlin-cognato et al., 2006; Hoffman et al., 2006; Phillips et al., 2009, 2011). This east/west split likely dates back to the glacial cycles prior to the last glacial maximum, 60 to 180 thousand years ago (Harlin-cognato et al., 2006; Phillips et al., 2011). Sexual size dimorphism is more pronounced in SSL (Krüger et al., 2014) and male reproductive skew in GSL is believed to be moderate (Pörschmann et al., 2010). If mating system variation is associated with sex-biased dispersal (Kokko & López-Sepulcre, 2006), the stronger sexual dimorphism in SSL would predict more pronounced male-biased dispersal.

We used whole-genome resequencing data to investigate patterns of sex-biased dispersal in GSL and SSL, leveraging information from across all four genomic compartments: MT, X, AUT, and Y. We employed three complementary approaches: (1) qualitative comparison of population structure across all four compartments; (2) quantitative estimation of reproductive skew and migration rates based on simple, equilibrium population genetic models; and (3) we further tested for isolation by distance (IBD) and isolation by environment (IBE) to assess whether the genomic consequences of sex-biased dispersal are fully explained by geographic distance or require the additional incorporation of sex-specific environmental barriers (Jeglinski et al., 2015; O’Corry-Crowe et al., 2006). When interpreting the results, we consider confounding biological processes including sex-biased mutation rates, demographic perturbation and linked selection, as well as the impact of bioinformatic choices.

## Materials and methods

### Population sampling

We obtained a total of 123 tissue samples from flipper clips that were collected from pups as part of previously published projects and under relevant permits (Aurioles-Gamboa et al., 2009; Hoffman et al., 2006; Orr et al., 2012; Wolf et al., 2008). GSL sampling was conducted under permit PC 001-03/04 (export permit No. 099/04 SPNG); SSL samples were collected under Marine Mammal Protection Act permit 782-1532-02 and according to ASM guidelines (American Society of Mammalogists Animal Care and Use Committee 1998). Samples span the core geographical ranges of both species and include 94 GSL (35 females, 59 males) and 29 SSL (8 females, 21 males) from eight and eleven breeding populations, respectively (**Figure 1B, Table S1, Table S2**).

### Y-chromosome assembly

As a reference Y-chromosome, we used the genome assembly of the closely related Californian sea lion (CSL) generated by the Vertebrate Genomes Project from a male sample sequenced at high coverage (GenBank: GCA_009762305.2; Peart et al., 2021). Because reliable variant discovery on Y chromosomes is challenged by extensive repetitive sequence, gametologous regions shared with the X chromosome and recent segmental duplications, we independently validated Y-linked genotype calls using a targeted genotyping assay. We designed 72 candidate Y-linked SNP assays using the MassARRAY® System by Agena Bioscience® (Ellis & Ong, 2017). Assay design and genotyping were performed at the Institute of Clinical Molecular Biology, Christian-Albrechts-University Kiel. After quality filtering, the final panel comprised 27 high-confidence, parsimony-informative Y-linked SNPs, which were successfully genotyped in 94 male GSL (**Supplementary Text 1, Table S3**). Genotype calls were then compared to bioinformatically inferred genotypes from whole-genome sequencing for the individuals common to both datasets (see results).

### Whole genome sequencing and data processing

For all 94 GSL and 29 SSL samples, we extracted genomic DNA using either the Monarch Genomic DNA Purification Kit or the Qiagen DNeasy Blood & Tissue kit according to the manufacturer’s protocols. Subsequent DNA quality checks were performed with Qubit, NanoDrop (A260/280, A260/230), and agarose gel electrophoresis. Sequencing libraries were prepared using the Illumina DNA Prep kit and sent to Novogene for sequencing on an Illumina HiSeq X (v2.5; 150 bp PE) for GSL, and a NovaSeq X Plus (150 bp PE; see **Table S2** for overview) for SSL. We obtained a mean autosomal coverage of 6.69 ± 1.11 for GSL and 8.56 ± 1.66 for SSL (**Table S4**). All individuals from a species were processed in the same way, so we do not expect batch effect within species.

For the GSL reads, which were generated on the HiSeq X platform, standard Illumina paired-end adapter sequences were trimmed at the first 13 bp (-paired --stringency 3 --length 40 *--a* and --*a2 “AGATCGGAAGAGC”)* and additionally with another detected adapter sequence (*--a “CACGTCTGAACTCCAGTCAC” --a2 “CGTCGTGTAGGGAAAGAGTG”*) using Trim Galore v0.4.1 (https://github.com/FelixKrueger/TrimGalore, a wrapper around Cutadapt v1.8.1 (Martin, 2011)) and FastQC v0.12.1 (Andrews, 2010). SSL reads, which were generated on the NovaSeq X Plus platform, were processed using the same tools and filtering criteria, with platform-specific adjustments: adapters were detected automatically by default, and polyA and polyG sequences were removed from both read ends (*-a A{10} -a2 A{10} -a G{10} -a2 G{10}*). Trimmed reads were then mapped to the male CSL reference assembly using BWA-MEM v0.7.17, with the option *-M* for Picard compatibility (Li & Durbin, 2009). Subsequently, the alignments were sorted, and the read groups were added using Picard v2.20.4 *SortSam* and *AddOrReplaceReadGroups*. For each individual, we merged lane-level BAM files and removed PCR/optical duplicates in Picard *MarkDuplicates* (Broad Institute, 2019). To improve alignment around indels, we performed local realignment using GATK v3.8 *RealignerTargetCreator* followed by *IndelRealigner*, with default settings (McKenna et al., 2010). We discarded secondary and supplementary alignments, unmapped reads, failed platform/vendor quality checks and PCR or optical duplicates using *samtools* with flag *–F 3844;* additionally, we filtered for paired alignments with low mapping quality with *–q 20*.

To identify and remove repetitive elements from AUT, X and non-pseudoautosomal region (non-PAR) of the Y, we used two repeat libraries. To create the first, we generated a *de novo* assembly-specific repeat library in RepeatModeler v2.0.5 (Flynn et al., 2020). From the obtained repeats, we excluded those overlapping with protein coding regions based on the UniProt database (The UniProt Consortium et al., 2025) using BLASTX v2.15.0 (Camacho et al., 2009) with the following settings: ≥85% sequence identity, high confidence (*-evalue 1e-5*), and minimum length (*-max_target_seqs 10*). For the second repeat library source, we used the published Dfam repeat library for eared seals (*Otariidae*). The two combined libraries were then used to annotate repetitive elements in the reference assembly with RepeatMasker v4.1.7-pl (Smit et al., 2015). All annotated repeats were excluded from downstream analyses.

In addition to repetitive elements, for all the compartments (AUT, X, Y, and MT), we identified an additional one-to-many sequence similarity to minimise misalignment effects. For that we compared each target chromosome with the rest of the genome using BLAST, and filtered out high-quality hits with ≥90% identity and a minimum length of 20 bp.

### Variant calling

For each genomic compartment, we performed variant discovery using *bcftools mpileup* v1.21 (Li & Durbin, 2009) with mapping and base-quality filters (*-q 30 -Q 30*). We then generated genotype calls by running *bcftools call* with *--ploidy 1* for haploid compartments (Y, MT), with a default ploidy of 2 for the AUT, and with a supporting ploidy file that included assigned sex for the heterogametic X. To ensure comparable statistical power and chromosomal properties among the X and AUT, variant calling for the autosomes was restricted to chromosome 9, which most closely matches the X chromosome in both length and GC content. We investigate the impact of this reduction in **Supplementary Text 2.1**. Variant sites overlapping annotated repeat intervals or homologous regions were excluded from downstream analyses using *bedtools subtract* v2.31.0. (Quinlan & Hall, 2010).

To obtain high-confidence SNPs, we applied a second stage of quality filters implemented in a custom Python script. For all compartments, we retained only sites that satisfied the following criteria: <10% missing genotypes across all individuals of a species; site QUAL and mapping quality MQ ≥60 for Y and ≥40 for the AUT, MT and X; read depth below the rounded product of the number of samples and the 85th percentile of the samples’ coverage distribution to exclude sites with excessive coverage; stricter <25% of genotypes exhibiting strand bias in a higher-coverage MT and more relaxed <50% of AUT, X and Y; variable sites in MT and Y supported by at least two samples differing from the consensus nucleotide; and <25% pseudo-heterozygous genotypes in MT and <50% in Y (**Table S4**). Altogether, this conservative filtering strategy reduced the risk of interpreting mapping artefacts or sequencing errors as true genetic variation, which is particularly important given the limited effective sequencing depth of the dataset. We identified likely parent-offspring or sibling pairs using the KING kinship coefficients implemented in PLINK2 v2.0.0-a.6 (Chang et al., 2015; Manichaikul et al., 2010) and excluded one GSL individual from a pair with a value exceeding 0.177, the recommended inference criterion for 1st-degree relationships (Manichaikul et al., 2010).

For haploid datasets, we additionally removed sites showing evidence for recombination based on the four-gamete test implemented in DNAsp v6.12.03 (Rozas et al., 2017). For the Y chromosome, missing genotypes were imputed using a random forest algorithm (RandomForestRegressor) implemented in the Python package scikit-learn v1.5.2 (Pedregosa et al., 2012). This approach was used to increase the number of callable sites while limiting the influence of stochastic missingness and sequencing noise.

### Analysis of population structure

Distinct populations or conservation units of marine mammals are referred to as stocks, and it is at this level we carried out analyses. We used both the scientific literature and the autosomal data to delimit the stock boundaries. For SSL, the literature consistently refers to an Asian, Western and Eastern stock (e.g. Baker et al., 2005; Harlin-cognato et al., 2006; Phillips et al., 2011). In GSL, the Western and Eastern stocks are consistently described, with the occasional addition of a Northern stock (e.g. Jeglinski et al., 2015; Wolf et al., 2008).

### AUT and X chromosome

To assess the population structure of AUT and X, we first applied another filtering step on the vcf file using PLINK v1.9.0 (Purcell et al., 2007): we filtered out missing genotype rates: missing call rates exceeding 5% (*--geno 0.05*) and missing sample rates exceeding 10% (*--mind 0.1*), and retained only variants with a minor allele frequency (MAF) ≥0.01 (*--maf 0.01*). To reduce the influence of linked markers, variants were pruned for linkage disequilibrium (LD) using PLINK. For PCA, LD pruning was performed with *--indep-pairwise 200 50 0.1*, whereas for ADMIXTURE analyses we used the more stringent parameters *--indep-pairwise 50 5 0.2*. We then conducted the PCA from the Python package scikit-learn v1.5.2 (Pedregosa et al., 2012) and performed admixture analysis using ADMIXTURE v1.3.0 (Alexander et al., 2009) with K-values from 2 to 5.

We also inferred spatially informed population structure in TESS3R v3.0 (Caye et al., 2016). To accommodate mixed-ploidy data, we first generated X and AUT pseudo-haploid calls in ANGSD v0.940 (Korneliussen et al., 2014) for the positions retained in the filtered dataset described above. We applied the GATK genotype likelihood model with allele frequencies as priors and generated pseudo-haploid genotypes by calling the most probable allele (-*sites -doPost 1 -gl 2 -dohaplocall 2*). We assessed whether this approach was consistent with the original variant-calling procedure in **Supplementary Text 2.2**. We subsequently ran TESS3R on the diploid AUT and pseudo-haploid X datasets using the least-squares algorithm, with the geographic regularisation parameter optimised by cross-validation across K values ranging from 2 to 9.

### MT and Y chromosome

For MT and Y, we produced haplotype networks using PopART v1.7 (Leigh & Bryant, 2015), using the median-joining approach (default ε = 0).

### Summary statistics of genetic variation within-and between-stocks

We used the Python library *allel* to compute F_ST_ between stocks using Hudson’s F_ST_ estimator, which is less sensitive to differences in sample size and mutation rate among compartments (Bhatia et al., 2013; Jong et al., 2024). For the haploid compartments, we additionally calculated Φ_ST_ using the *ape* and *pegas* R packages (Paradis, 2010; Paradis & Schliep, 2019). For X and AUT, we also computed Watterson’s θ in females using ANGSD in windows of 50kb with 10kb steps and with minimal filtering: a minimum depth of 2 (*-setMinDepth*), maximum 10% of missing data (*-minInd*) and a minimum mapping quality (*-minMapQ*) of 40 (**Supplementary Text 2.3**).

### Isolation-by-distance (IBD) and isolation-by-environment (IBE)

For IBD and IBE analysis, we used the following genetic, geographic and environmental distance matrices. Pairwise genetic distances were based on linearised Hudson’s *F_ST_* (Rousset, 1997) for AUT and X-chromosomal datasets, and Φ*_ST_*for haploid MT and Y-chromosomal datasets. Pairwise geographic distances between rookeries were calculated based on GEBCO bathymetry data (GEBCO Bathymetric Compilation Group, 2025). We calculated coastal, depth-weighted pairwise distances between rookeries as least-cost paths across a bathymetric raster projected in UTM coordinates (metres). Land and no-data cells were treated as impassable, whereas ocean cells were assigned depth-dependent costs to favour routes along continental shelf areas. Rookery coordinates (WGS84) were projected onto the nearest water pixels, and the minimum accumulated cost distance between each pair of rookeries was calculated using the MCP Geometric function from the Python scikit-image package (Van Der Walt et al., 2014). Distances were converted to kilometres to yield the final pairwise symmetric matrix. Environmental distances were represented as discrete groupings based on ecological contrasts described in the literature, and supported by visual assessment of key environmental variables, including sea surface temperature (average over 8 years between 2002 & 2010, “MS_biogeo13_sst_mean_5m”), bathymetry (“MS_bathy_5m”) and primary productivity (average over 14 years between 2000 & 2014, “BO21_ppmean_bdmean”) obtained and plotted using the R packages sdmpredictors, terra and raster (Bosch & Fernandez, 2023; Hijmans, 2025a, 2025b) (**Figure S3**). In GSL, rookeries of the Western stock (FH and IB) were classified as environmentally distinct from the rest of the populations, as they benefit from higher productivity and colder waters (**Figure S3**) (Jeglinski et al., 2015). In SSL, Eastern rookeries (GRGR, HAZY, TRIA, and WSIS) formed a separate group from the rest, as they experience warmer waters (**Figure S3**).

We then applied Mantel and partial Mantel tests implemented in the R package *vegan* (Oksanen et al., 2025) to assess associations between genetic, geographic and environmental distance matrices. Following Shafer and Wolf (2013) we conducted five Mantel tests: (1) genetic versus geographic distances, (2) genetic distances versus environmental dissimilarity, (3) geographic distances versus environmental dissimilarity, (4) a partial test correlating genetic distance versus geographic distance while controlling for environmental dissimilarity, and (5) a partial test correlating genetic distance versus environmental dissimilarity while controlling for geographic distance. In addition, we applied Multiple Matrix Regression with Randomisation (MMRR) using the R package *ecodist* (Goslee, 2010) to assess covariance between distance matrices. For both Mantel and MMRR analyses, we evaluated statistical significance with permutation tests (9,999 permutations). For the IBD plots, we set all negative genetic distances to zero, as they suggest no differentiation between groups and lack biological interpretation.

### Inference of sex-biased dispersal

We estimated sex-biased migration rates under migration–drift equilibrium within an island model framework. Details of the inference are provided in **Supplementary Text 3 & 4**. In brief, we first estimated sex-specific effective population size (*Ne^M^* and *Ne^F^)* as a function of nucleotide diversity from autosomal (θ *= 4Ne*μ) and X-linked (θ *^X^ = 4Ne ^X^*μ*^X^*) compartments, accounting for male-bias in mutation rate (α), with α set to 3 (Amster & Sella, 2016; Foote et al., 2016). We then inferred sex-specific migration rates (*m^M^*, *m^F^*) by exploiting sex-specific *Ne,* as well as inheritance patterns of genomic compartments, and their corresponding pairwise differentiation (Hudson’s *F_ST_*) following the rationale outlined in Hedrick et al. (2015). Those estimates integrate both the sex-specific migration rates (*m^M^* and *m^F^*) and the corresponding effective population sizes (*Ne^M^* and *Ne^F^*), which are expected to differ between males and females in a polygynous system. Direct estimates were obtained from haploid, uniparentally inherited markers, where differentiation of the Y chromosome and mitochondrial DNA provides information on male- and female-mediated gene flow, respectively. Indirect estimates were obtained by combining autosomal vs mitochondrial and autosomal vs X chromosomal differentiation.

## Results

### Genetic variation of the Y chromosome

The male-specific region of the Y chromosome assembly, excluding the pseudoautosomal region, spans 3,986,568 bp (**Figure 2, Figure S4**). We identified 55 gene models, including 18 pseudogenes, 7 multi-copy genes and 30 single-copy genes, including 4 gametologs. All single-copy genes but one (TCEANC) had sufficient sequence divergence to reliably distinguish the X- and Y-linked copies using a 95% sequence identity threshold. However, shorter regions of X-Y similarity, including partial gene sequences and non-coding regions, can still lead to read mismapping. Low-complexity repetitive sequences accounted for more than half of the assembly (∼57%). Following stringent filtering to retain only sites unambiguously assigned to the Y chromosome, we used 1,436,828 bp (36% of the male-specific region) for population genetic analyses. Across these regions, we identified 50 and 49 SNPs conforming to our quality criteria and segregating in the GSL and SSL populations, respectively (see methods). These variants were predominantly located within single-copy genes (40% and 59%, respectively). Genotype validation of a subset of 15 SNPs segregating in 62 GSL individuals and overlapping with the MassARRAY set of genotypes showed 100% concordance, 12 SNPs did not intersect with the MassARRAY (see **Table S3)**.

**Figure 2.**
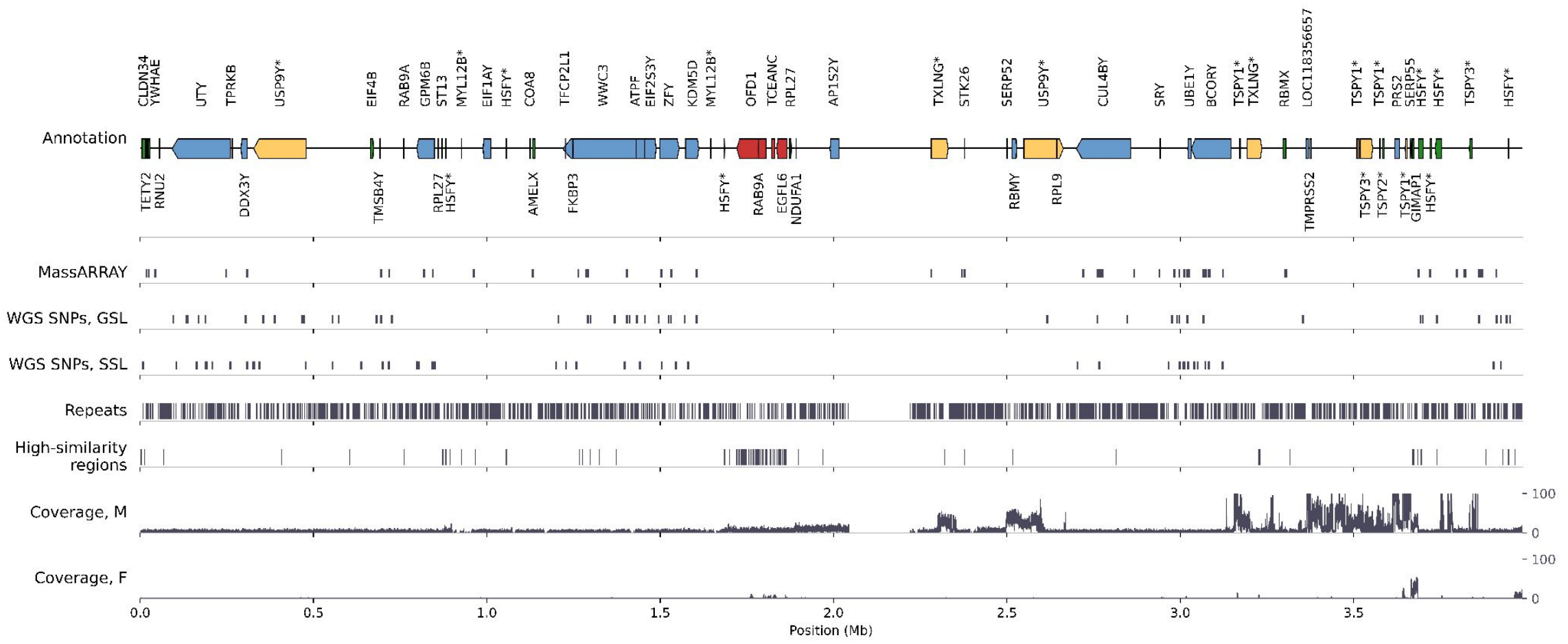
Characterisation of the Y chromosome as a potential resource for population genetic analyses. The schematic represents the 3,986,568 bp male-specific region of the Y chromosome from the Californian sea lion reference genome assembly used in this study, excluding the pseudoautosomal region. The genomic tracks are displayed from top to bottom as follows: i) Gene models (including exons and introns) are represented by arrows indicating strand orientation (+, rightward; −, leftward). Colours indicate gene categories, including putative single-copy Y genes (blue), multi-copy Y genes (yellow & asterisk in gene name), pseudogenes (green), _and_ gametologs with high similarity to the X (red). ii) SNPs from an independent genotype assay (MassARRAY) and those remaining after filtering the WGS data in GSL and SSL for iii) repeats, iv) regions with high sequence similarity to the rest of the genome and coverage for v) one male (FH01) and vi) one female (EG05) GSL.

### Sex-biased migration

In total, we inferred 143,765 and 144,265 variants segregating in 93 GSL and 29 SSL individuals, respectively, and distributed across the four genomic compartments: MT (GSL: 57 & SSL: 50), X (30,928 & 27,495), AUT (112,730 & 116,707) and Y (50 & 49) (**Table S4**). Based on these data, we explored sex-biased migration by comparing population structure, fixation indices and derived statistics, as well as patterns of IBD and IBE across genomic compartments. Results are presented for two stocks in GSL and three stocks in SSL, following previous literature and the genetic structure inferred from the autosomal data (**Figure S5**). In the Supplement, we additionally explore finer population structure in GSL by subdividing the Eastern stock into Northern (PC, GE) and Central (remaining Eastern rookeries) groups (**Figure S6** & **S7**). For SSL, we examine the effect of merging the Asian and Western stocks into a single group (**Figure S8 & S9**).

### Galápagos sea lion

Population structure in GSL was overall weak for X and AUT, with each PC axis explaining <3% of the variance. Nevertheless, PCA, admixture analyses and spatial genetic clustering revealed clear differentiation between the Western (FH, IB) and Eastern stock (remaining rookeries) (**Figure 3A** right, **Figure S6, Figure S7**). Contrary to the expectation from *Ne* and male-biased dispersal, population stratification appeared slightly stronger for AUT than for the X chromosomes (AUT: 15% vs X: 27% mixed-ancestry individuals), even when subsampling the AUT data to the same number of variants as the X (**Supplementary Text 2.4**). However, consistent with male-biased gene flow, stock structure was slightly more pronounced in the mtDNA than in the Y haplotype network (**Figure 3A** left). Haplotype diversity and stock-specific proportions of haplotype groups (private haplotypes) were broadly similar between MT and Y, providing no conclusive information on sex-biased gene flow (**Table S5**).

**Figure 3.**
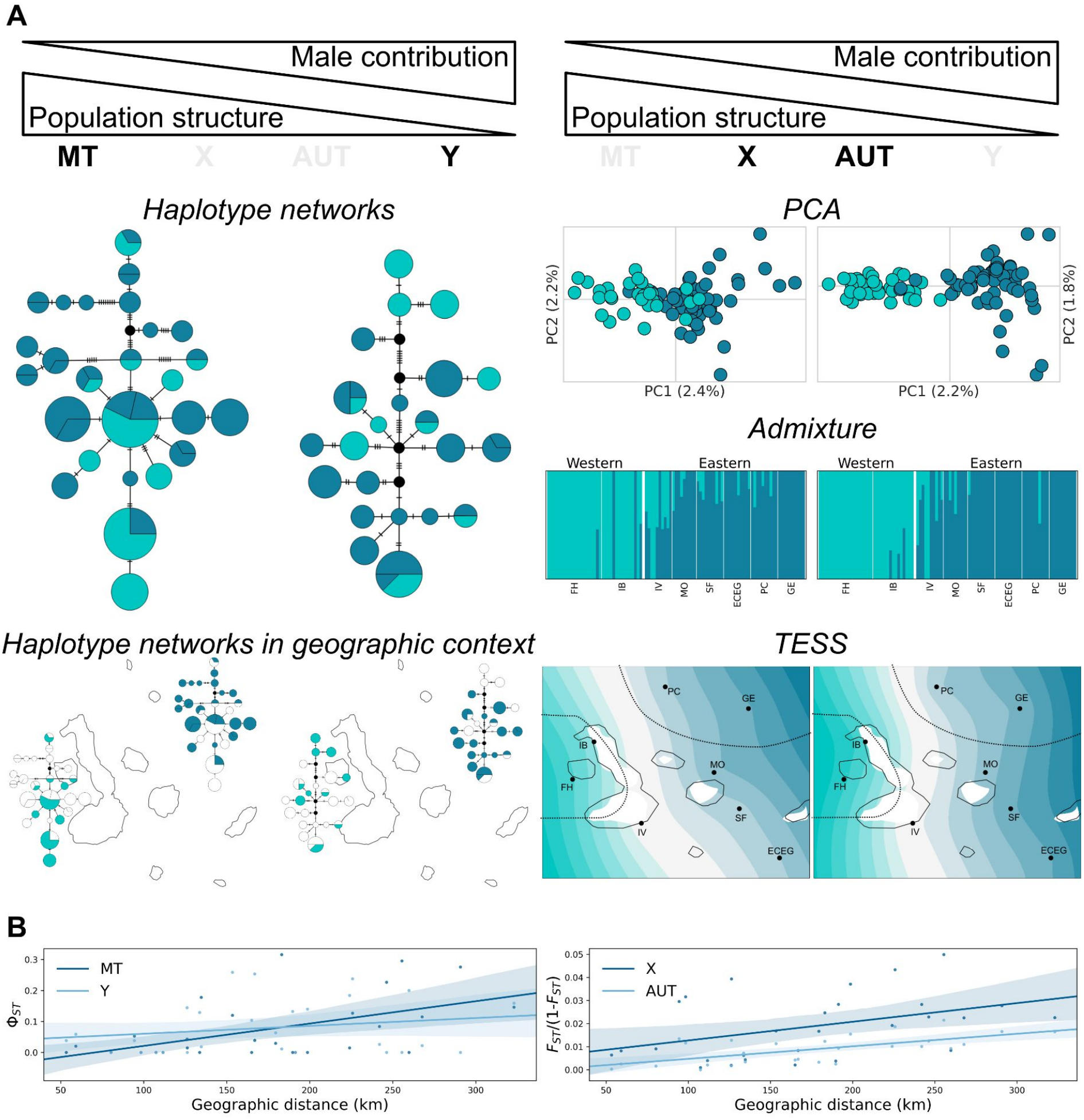
Population structure across genomic compartments in GSL. A) Visualisation of the population structure of the two main stocks shown for haploid (*left:* MT, Y) and diploid (*right:* X, AUT) genomic compartments. Colours represent stocks as defined in the literature for haplotypes and PCA, and as assigned by the analysis in TESS and admixture (Western stock: green, Eastern stock: blue). The top row shows the male contribution to genetic variation in each genomic compartment and the expected effect on population subdivision. *Left:* Haplotype networks for MT and Y showing haplotypes from both stocks jointly (*above*) and separately in a geographic context (*below*). *Right:* PCA (*top*), admixture proportions (*centre*), and TESS geographic interpolation with rookery abbreviations, where the dashed line separates stocks (*bottom*). B) Isolation-by-distance (IBD) calculated as linearised F_ST_ for diploid compartments (*right*) and Ф_ST_ for haploid compartments (*left*) vs geographical distance, with linear regression and 95 % CI.

To obtain a quantitative estimate of population differentiation, we calculated *F_ST_* between rookeries and stocks for each genomic compartment. Differentiation of rookeries from distinct stocks consistently exceeded that observed within the same stocks (**Table S6, Table S7**), and differentiation between stocks followed expectations for male-biased dispersal (*Φ_ST_*: MT=0.149 > Y=0.058; *F_ST_*: X=0.015 > AUT=0.009). To quantify sex-bias in gene flow between stocks, we next calculated the ratio *Ne^F^/Ne^M^,* which we later integrated into the *m^M^*/*m^F^* computation. Considering the polygynous mating system, we do not assume equal variance in reproductive success between males and females and need to control for differences in male (*Ne^M^*) and female (*Ne^F^*) effective population sizes to obtain an independent estimate of male-biased migration *m^M^* / *m^F^*. Consistent with moderate polygyny in GSL, we inferred an elevated number of effectively breeding females (*Ne^F^/Ne^M^* of 1.284; **Table S8**). We then estimated *m^M^*/*m^F^*, assuming a male-mutation bias of α=3 (see methods and **Supplementary Text 4**). This yielded an estimate of up to sevenfold higher migration in males relative to females (*m^M^* / *m^F^*) (**Table 1, Table S9**).

**Table 1:** The ratio of migration rates for males (*m^M^)* and females (*m^F^*) as estimated from inter-stock F_ST_ values for MT and AUT and from ratios of effective population sizes (*Ne^F^/Ne^M^*) reflecting differences in reproductive skew, estimated from genetic variation in X and AUT. Stock definitions used in the main analyses are shown in black, whereas additional population subdivisions explored in the Supplement are shown in grey.

| **Species** | **Nr. stocks** | **Stock pairs** | **Ne<sup>F</sup> / Ne<sup>M</sup>** | **m<sup>M</sup> / m<sup>F</sup>** |
|---|---|---|---|---|
| GSL | 2 | Western - Eastern | 1.285 | 7.045 |
|  | 3 | Western - Central |  | 6.268 |
|  |  | Western - Northern |  | 6.844 |
|  |  | Central - Northern |  | 1.803 |
| SSL | 2 | Eastern – Asian/Western | 7.016 | 42.377 |
|  | 3 | Asian - Western |  | 16.119 |
|  |  | Asian - Eastern |  | 28.217 |
|  |  | Western - Eastern |  | 72.426 |

Finally, we investigated genetic differentiation across genomic compartments as a function of geographic distance (IBD) and environmental distance (IBE) (**Figure 3B**). Consistent with the expectation of male-biased dispersal, the Y chromosome showed no significant pattern of IBD, whereas significant IBD was detected for AUT, X and MT (**Table 2**, for multiple matrix regression results, see **Table S10).** After controlling for environmental variation using partial Mantel tests, the signal remained significant only for the autosomes (**Figure S10**). MT further showed a strong and significant pattern of IBE after controlling for geographic distance.

**Table 2:** Correlation coefficients of Mantel tests for isolation by distance (IBD) and isolation by environment (IBE). Genetic distance (GenD) was measured as linearised F_ST_ for diploid compartments and Ф_ST_ for haploid compartments, geographic distance (GeoD) as the pairwise swimming distance between rookeries (km), and environment (Env) as the ecological grouping variable. Statistically significant results (P<0.05) are shown in bold.

| **Species** | **Compartment** | **GenD ~ GeoD** | **GenD ~ Env** | **GenD ~ GeoD \| Env** | **GenD ~ Env \| GeoD** |
|---|---|---|---|---|---|
| GSL | MT | **0.521** | **0.719** | 0.259 | **0.617** |
|  | X | **0.419** | 0.29 | 0.329 | 0.099 |
|  | AUT | **0.648** | 0.446 | **0.547** | 0.177 |
|  | Y | 0.208 | 0.165 | 0.146 | 0.07 |
| SSL | MT | **0.345** | **0.375** | **0.327** | 0.359 |
|  | X | **0.602** | 0.281 | **0.597** | 0.266 |
|  | AUT | **0.521** | **0.343** | **0.516** | 0.333 |
|  | Y | -0.06 | 0.011 | -0.062 | 0.018 |

Overall, these results are qualitatively consistent with a moderately polygynous mating system associated with elevated male-biased migration reducing population structure in the Y chromosome relative to mtDNA. Although differences between the autosomes and the X chromosome did not always conform to the expectation, the corresponding estimates of the male-to-female migration ratio (*m^M^/m^F^*) similarly indicated elevated male migration.

### Steller sea lion

In the Steller sea lion, PC1 of both X and AUT separated the Eastern stock from both Western and Asian stocks, which were incompletely separated along the PC2 axis (**Figure 4A** right). Consistent with the expectation from *Ne* and male-biased dispersal, population structure was slightly stronger for X than for AUT (PC1: 7.1% vs. 6.4%). In contrast to the PCA, admixture analysis showed more admixture of Eastern and Western individuals than for the Asian stock, with generally elevated admixture in the X (X: 45% vs. AUT: 31% mixed-ancestry individuals) (see **Figures S8 & S9** for the Asian and Western stocks collapsed). This pattern remained after sub-sampling AUT SNPs to the same number of variants as the X (**Supplementary Text 2.4**). Spatial-genetic clustering portrayed a narrower Western stock representing only a single rookery in X (BULD), and comprising two rookeries for AUT (BULD & WALR; see **Figure S9** for merged stocks). Haplotype networks revealed stronger stock structure (**Figure 4A** left) and a higher proportion of stock-specific haplotypes for MT than for Y, but these patterns alone do not establish sex-biased gene flow (**Table S5**).

**Figure 4.**
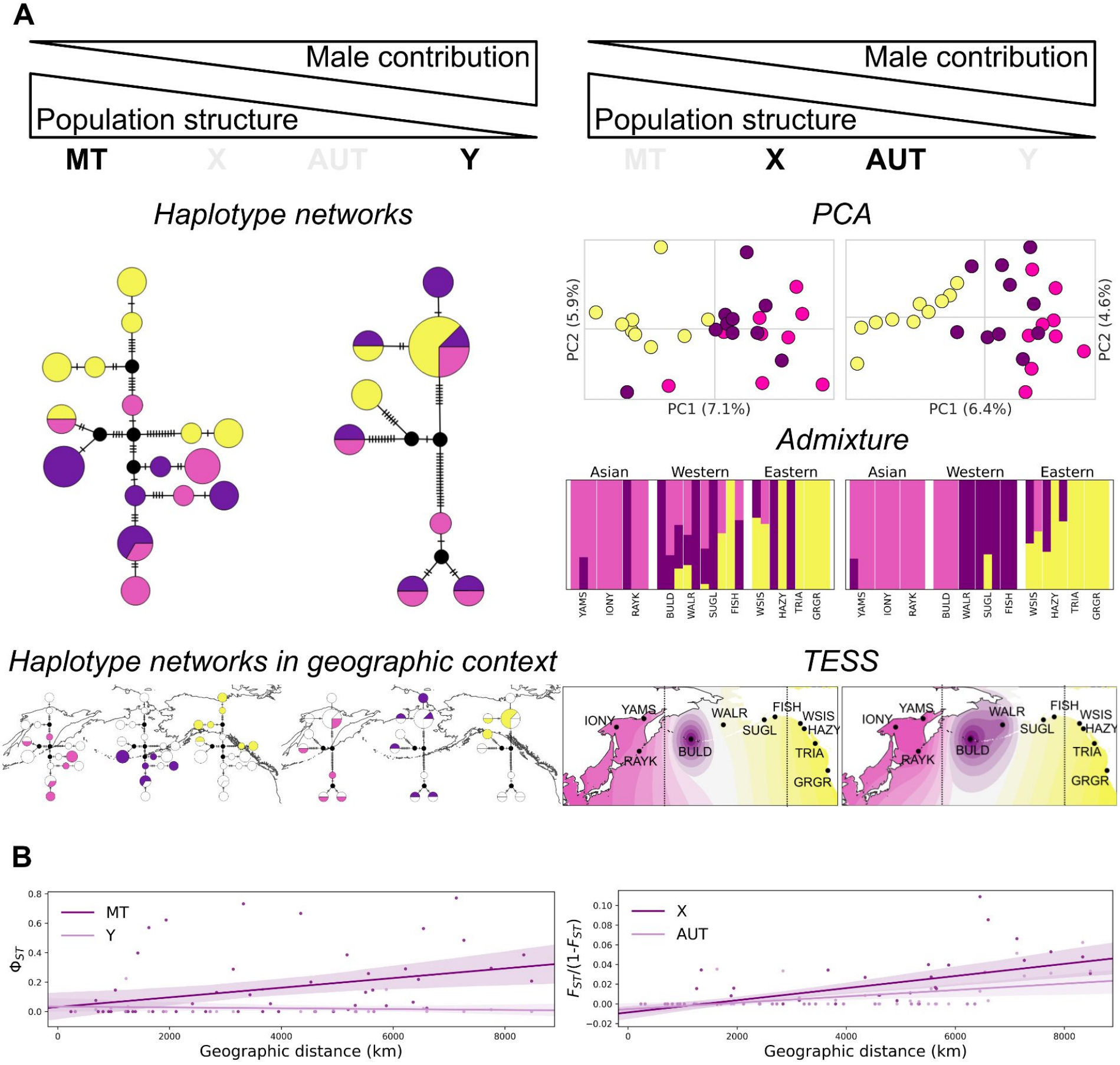
Population structure across genomic compartments in SSL. A) Visualization of population structure of the three stocks shown for haploid (*left:* MT, Y) and diploid (*right:* X, AUT) genomic compartments. Colours represent stocks as defined in the literature for haplotypes and PCA and as assigned by the analysis in TESS and admixture (Asian stock: pink, Western stock: purple, Eastern stock: yellow). The top row shows the male contribution to genetic variation in each genomic compartment and the expected effect on population subdivision. *Left:* Haplotype networks for MT and Y showing haplotypes from both stocks jointly (*above*) and separately in a geographic context (*below*). *Right:* PCA (*top*), admixture proportions (*centre*), and TESS geographic interpolation with rookery abbreviation and where the dashed line separates stocks (*bottom*). B) Isolation-by-distance (IBD) calculated as Ф_ST_ vs. geographical distance for haploid compartments (*left*) and diploid compartments (*right*) with linear regression and 95 % CI.

As in GSL, pairwise F_ST_ between all populations and the two major stocks showed differences between genomic compartments consistent with male-biased migration (*Φ_ST_*: MT=0.387 > Y=0.022; *F_ST_*: X=0.032 > AUT=0.021, see **Table S6, Table S7** for all three stocks). SSL showed a more pronounced bias in the number of effectively breeding females (*Ne^F^/Ne^M^* = 7.016; **Table 1**, **Table S8**), consistent with strong reproductive skew in males. Overall, after accounting for sex-specific *Ne,* this resulted in substantially higher estimates of male-biased migration (*m^M^* / *m^F^*) than in the GSL, ranging from 16 – 72 **(Table 1, Table S9**).

Genetic differentiation for MT, X and AUT showed a significant association with geographical distances, but not with the environment (**Figure 4B**, **Table 2**). This effect remained after controlling for the environment (**Table S10, Figure S10)**. As in GSL, the Y chromosome showed neither significant IBD nor IBE associations.

Overall, these results provide concordant indication for both stronger polygyny and male-mediated gene flow among stocks in SSL than in GSL.

## Discussion

In this study, we took a genomic approach to investigate mating systems and sex-bias in dispersal behaviour in two highly mobile, polygynous marine mammal species. Our approach was based on the assumption that these behaviours leave predictable signatures in genetic diversity within and among populations across genomic compartments with contrasting modes of sex-specific inheritance. In addition to the commonly used female-inherited mitochondrial genome, and the biparentally transmitted autosomal markers, we included both the shared (X) and the male-limited (Y) chromosomes. To date, whole-genome sequencing of the latter has mostly been explored in humans (Lippold et al., 2014; Sayres et al., 2014), but is still rarely applied elsewhere (de Jong et al., 2025; Felkel et al., 2019; Laso-Jadart et al., 2025; Smeds et al., 2019). We used differences in genetic diversity among these compartments to infer sex-specific reproductive skew and migration as proxies for mating system and dispersal, adopting a simple mutation–migration–drift equilibrium model. This approach comes with several important limitations that are rarely made explicit. Below, we first interpret the observed patterns at face value in light of the species’ biology and then consider potentially confounding processes and methodological challenges that may influence interpretation of the results.

### Reproductive skew in males

In both species, male *N*e was reduced relative to female *N*e as expected for higher reproductive skew among males and the polygynous mating system. However, this reduction was substantially greater in SSL than in GSL, consistent with the greater sexual size dimorphism in SSL (3.48 vs. 2.56, Lindenfors et al., 2002). Sexual size dimorphism is generally associated with stronger sexual selection and intensity of competition among males in polygynous pinnipeds, but it is not a perfect predictor of reproductive skew (Krüger et al., 2014; Lindenfors et al., 2002). Sexual skew may be mediated by ecological factors. For instance, the unusually long breeding season in the tropical GSL (∼5.2 months) constrains the ability of territorial males to monopolise access to receptive females, making colony attendance a key determinant of male reproductive success (Krüger et al., 2021; Pörschmann et al., 2010). Although territorial males achieve the highest reproductive success per individual, only 5–10% of males are territorial, and non-territorial males sire more than 80% of pups for which paternity could be assigned each year (Meise et al., 2014; Wolf et al., 2005). In contrast, reproduction in SSL is more synchronised and concentrated over a substantially shorter seasonal window (∼3.8 months) facilitating the effective monopolisation of females by territorial males (Gentry, 1970).

### Male-bias in dispersal

Theory suggests that polygynous mating systems may also be prone to male-biased dispersal (Greenwood, 1980; Li & Kokko, 2019). Our estimates support this idea. Male bias in migration was inferred for both species and was more pronounced in the SSL, where reproductive skew was also greater. In the following, we discuss the migration patterns of both species in the context of the prevailing environmental conditions and their contrasting life histories.

GSL dispersal ability largely exceeds their small geographical range as they can cross the species range within a few days (Jeglinski et al., 2015) and, in exceptional occurrences, have been recorded as far as Mexico (Ceballos et al., 2010; Elorriaga-Verplancken et al., 2022). In this species, population structure can therefore only arise through substantial breeding site or habitat fidelity. Population structure was shallow, but most strongly marked between Western and Eastern stocks (as seen in Peart et al., 2020), broadly aligning with a marked ecological gradient across the archipelago that has previously been discussed in the context of dietary adaptation (Wolf et al., 2008). The male-biased migration rate inferred in this study, combined with the strong IBE signal in MT, suggests that population structure is influenced by habitat preferences of females. While GSL females certainly benefit from familiarity with their environment (Drago et al., 2016), prolonged lactation and simultaneous care for offspring of different ages further limits the dispersal of adult females (Childs et al., 2025; Kalberer et al., 2018; Krüger et al., 2021; Maniscalco, 2014; Trillmich & Wolf, 2008; Trites, 2021). But even prior to sexual maturity, persistent mother-daughter associations and a tendency of female pups to settle close to their mothers in a familiar social environment may further reinforce female philopatry (Wolf et al., 2007; Wolf & Trillmich, 2007).

In SSL, female dispersal appears less restricted, as weaning generally occurs before one year of age and, in contrast to GSL, SSL females rarely breed while caring for a juvenile from the previous year (Hastings et al., 2017; Raum-Suryan et al., 2002; Trites, 2021). Accordingly, mitochondrial differentiation remained low between the Western and Asian stocks, suggesting females disperse between regional rookeries across much larger geographic distances than GSL. Nevertheless, familiarity with the environment, territorial breeding males, and neighbouring females still appear to restrict female dispersal (Hastings et al., 2017). Although we found no evidence of female-specific IBE as in GSL, our estimates suggest that the sea surface temperature boundary between the Eastern and Western stocks may restrict female-mediated gene flow (**Figure S3**). Male migration rates were an order of magnitude higher than female migration rates, suggesting substantially fewer constraints on male dispersal, which aligns with the idea that the strongly polygynous mating systems promote long-distance movements of adult males along coastal routes in search of mating opportunities (Maniscalco, 2014; Sweeney et al., 2025).

### Limitations

Overall, our inference from genomic data appears to align with the available behavioural, environmental and life-history information and expected dispersal patterns in both species. However, sex-specific estimates should be interpreted with caution, as they are influenced by confounding biological factors, as well as by bioinformatic choices, that we sought to minimise but could not fully eliminate.

### Confounding factors

Genetic diversity among genomic compartments is differentially affected by multiple evolutionary processes, which in turn influence estimates of reproductive skew and sex-biased migration. We will discuss each of these in turn.

### Sex-specific mutation rates

Males and females differ in the rate of germline mutation. Sperm production involves more cell divisions, and thus more potential errors in replications (i.e. mutations) (Ellegren, 2007; Miyata et al., 1987; Wilson Sayres, 2018), generally increasing with paternal age (Amster et al., 2020; Amster & Sella, 2020). Because autosomes and X chromosomes differ in their transmission through males and females, they experience different contributions from sex-specific mutation rates. This pattern can be exploited to infer sex-bias in mutation rate e.g. via comparison of neutral substitution rates between X and AUT along deeply diverged lineages (Miyata et al., 1987).

In the context of this study, sex-biased mutation rates can alter genetic diversity on the X chromosome relative to the expected three-quarter value of autosomal diversity under a Wright-Fisher model (Corl & Ellegren, 2012). This matters, since estimates of male (*Ne^M^*) and female (*Ne^F^*) reproductive skew, themselves affecting sex-biased migration rates (*m^M^*, *m^F^*), are derived from the effective population sizes of AUT and X, which thus crucially depend on the value of male-bias in mutation α (**Supplementary Text 3**). We therefore conducted sensitivity analyses for the published range of α from 2 to 3 (Amster & Sella, 2016; Foote et al., 2016). Qualitatively, male-reproductive skew (*Ne^F^/ Ne^M^* >1) was stably inferred across most of the parameter space, but values could differ by up to 1.46 -1.75-fold for GSL and SSL, respectively (**Table S8,** see also **Figure S1**). To reduce this uncertainty, future research would benefit from species-specific pedigree-based estimates of mutation rates for both sexes in the target species (Bergeron et al., 2023).

Mutation rates of the mitochondrial and Y chromosomes are known to differ substantially from one another and from those of the autosomes and X chromosome (Melvin & Ballard, 2017; Sayres et al., 2014). Nevertheless, in the context of this study, MT and Y are still informative for tracing female and male migration. Assuming that migration rates are substantially greater than mutation rates and that *F*-statistics are mostly sensitive to the former (*F_ST_* ≈ 1/[1+4*Ne*(*m*+μ)]), mutation rates of those compartments can likely be neglected. Importantly, our estimates of *Ne^F^/ Ne^M^* were based on AUT and X, not on MT and Y (see above, **Supplementary Text 3**).

### Demographic history

Population size changes can also affect genomic compartments differently, which complicates the interpretation of relative diversity further. For example, X-linked nucleotide diversity is more strongly reduced than AUT upon population size reduction, but is also faster to recover (Pool & Nielsen, 2007). The magnitude of the reduction, as well as its duration and timing, also interact with the breeding sex ratio, the temporal dimension of (sex-specific) migration and sex-specific survival, resulting in complex patterns of ancestry and genetic diversity across AUT, X, Y and MT (Goldberg & Rosenberg, 2015; Miyagi et al., 2025; Spurley & Payseur, 2025), as observed, for example, between Neanderthals and modern humans (Platt et al., 2026).

Our estimates of *Ne^F^/ Ne^M^* derived from X and AUT are based on the population mutation parameter θ under the assumption of mutation–drift equilibrium in a stable Wright–Fisher population. This assumption is clearly violated in natural populations, including those studied here. Previous work suggests that both species, with effective population size estimates of ∼12,000-17,000, have experienced population size reductions to ∼1/3 of the ancestral population size (*2Ne*_anc_) over the duration of > 25,000-35,000 generations (Peart et al., 2020). To obtain an intuition about the magnitude of this effect on *Ne^F^/ Ne^M^*, we simulated the approximate demographic scenario following (Pool & Nielsen, 2007) (**Figure S2**). The model suggests that the X/AUT ratio may be downward-biased relative to equilibrium expectations (e.g. 0.625 for α=3) in both species, slightly more so in SSL than in GSL (**Figure S2A & B**). Since our estimates of *Ne^F^/Ne^M^* attribute variation in the X/AUT ratio exclusively to different operational ratios of the sexes, adjusting for the demographic effect would increase the X/AUT ratio, and with it the estimates of both *Ne^F^/Ne^M^*and *m^M^/m^F^* (**Figure S11**). As a consequence, both species may exhibit a higher degree of polygyny and a stronger male bias in dispersal, and the difference from SSL would be even more pronounced. To try to better accommodate demographic history, we focused our considerations on θ*^W^*rather than π, as it places greater weight on rare, younger variants and may be more sensitive to demographic differences and may at least mitigate this effect (for comparison, see below).

### Linked selection

Linked selection is arguably the most difficult confounding factor to quantify and control for, as it depends not only on the recombination rate and the distribution of fitness effects (Botero-Castro & Wolf, 2026), but also on the dominance of mutations (Hammer et al., 2010). Both uniparental compartments consist of haploid, non-recombining haplotypes, making them particularly susceptible to linked selection from mutations of any dominance coefficient. Moreover, gene content of the MT differs substantially from the Y with MT encoding OXPHOS genes that are subject to strong purifying selection (Nabholz et al., 2013). The single-locus nature, strong linked selection, and substantial genetic drift associated with their small effective population sizes make direct comparisons with the Y chromosome challenging. Nevertheless, despite the large expected variance introduced by the above processes, our study still seems to detect a signal that is consistent with the expected mode of dispersal.

Expected differences in genetic diversity that are due to linked selection also apply to X and AUT, which crucially affect our *Ne^F^/Ne^M^*and *m^M^/m^F^* estimates. In pinnipeds, the X chromosome is approximately average in size relative to the autosomes (Peart et al., 2021) (e.g., CSL: X 129 Mb vs AUT average 133 Mb), but we expect the X to exhibit lower recombination rates than AUT. This is due to the fact that the recombination rate of the X chromosome (r^X^) exclusively derives from females (r^X^ = 2/3 r^F^), whereas autosomes are equally affected by the recombination rate of both sexes (r^AUT^ = 1/2(r^F^+ r^M^)). This leads to a ratio of r^X^/r^AUT=^ 4r^F^/3(r^F^ + r^M^) which is lower than one even if female recombination is substantially elevated over that in males, as e.g. in humans by a factor of ∼1.6 (Kong et al., 2002). Moreover, in hemizygous males, (partly) recessive alleles are directly exposed to both positive and purifying selection on the X, while they are sheltered in a heterozygous state in AUT. As a consequence, linked selection, through both hitchhiking and background selection, is expected to reduce diversity on the X more strongly than on AUT (Begun & Whitley, 2000; Charlesworth et al., 1987; Hammer et al., 2008). The relatively stronger reduction of X-linked than AUT diversity due to linked selection is thus expected to affect the *Ne^F^/Ne^M^* in the same direction as population size reduction discussed above, increasing both the strength of polygyny and the extent of male-biased dispersal relative to our estimates (**Figure S11**). Although we do not know the magnitude of this effect, this thought experiment suggests that compartment-specific effects of linked selection may represent a substantial confounding factor. Incorporating this bias in analyses, however, is complex, as it requires thorough estimates of recombination rates and the distribution of fitness effects for each population and genomic compartment, which are already complicated for autosomal data (Botero-Castro & Wolf, 2026) and currently not available for pinnipeds and most other ecological and behavioural model species.

## Methodological considerations

While evolutionary processes contributing to changes in genetic variation have been studied extensively, technical effects have received less attention. Below, we explore the influence of methodological choices on our results and show that some analyses are highly sensitive to bioinformatic decisions.

### Comparability

One obvious methodological limitation is that analytical approaches are specific to particular genomic compartments, limiting comparability between haploid, uniparentally inherited genomes and the diploid autosomes or hemizygous X. PCA and admixture analyses require multi-locus data, and so are inherently unsuited to mtDNA and Y. Conversely, haplotypenetworks are straightforward to construct for mtDNA and Y, but require haplotype resolution, which is more difficult to infer for autosomes and the X. We therefore discuss the two groups of compartments separately.

### MT and Y chromosome

The comparison of MT and Y provides the most direct contrast between female and male genealogical histories. Yet, inference is limited, as the MT and Y each represent a single genealogy and are therefore sensitive to stochastic variation of the coalescent process. Additionally, the Y chromosome is particularly challenging to analyse because it exhibits high repetitiveness, duplicated genes and regions with high similarity to the rest of the genome. In this study, only 36% of Y chromosome data remained usable after filtering for repeats, high-similarity homologous regions and low-quality sites (see methods; **Figure 2, Figure S4**). Notably, 0.04% of the chromosome Y showed coverage >10,000x, consistent with the collapse of low-complexity regions during assembly (**Figure S4**). Collapse of the assembly, despite state-of-the-art methodology, is consistent with cytogenetic analyses of the reference genome (Peart et al., 2021) which show a Y chromosome moderately smaller than chromosome 17, whereas the assembly is substantially reduced (Y without PAR: ∼4 Mbp vs chr17: ∼60 Mbp). Our Y chromosome data are further subject to lower coverage and fewer individuals than all other compartments, affecting the relative confidence that can be placed in estimates of genetic variation.

For the reasons above, quantitative estimates of sex-biased migration rates directly based on *F_ST_^Y^* and *F_ST_^MT^* should be treated with due caution. Qualitatively, population structure and estimates of sex-biased migration still suggested male bias in SSL, but not in GSL (**Table S9, Supplementary Text 4**).

### X vs AUT

X and AUT are parental markers that provide multiple independent loci for inferring genealogies and deriving summary statistics for intra-and inter-specific genetic variation (e.g. Dahms & Momigliano, 2026; Platt et al., 2026; Spurley & Payseur, 2025). Compared to Y, the inference based on MT and AUT should therefore yield more reliable results, and when applied to our data show a robust male-biased migration signal for both SSL and GSL (**Table 1**, indirect ratios AUT-MT in **Table S9**). In principle, quantitative, chromosome-wide estimates from X and AUT should be the gold standard for sex-biased dispersal estimation. When applied to our data, it confirmed male-biased migration in GSL, but for SSL the observed combination of estimates was incompatible with the model’s parameter constraints; therefore, the results cannot be interpreted (indirect ratios AUT-X in **Table S9**).

A key methodological choice was comparing X-chromosome data to a single autosome matched for GC content and SNP number as closely as possible. We show that reducing the autosomal data to a single representative chromosome is appropriate here, and the results differ only slightly from when all autosomes are analysed together (**Supplementary Text 2.1**). Despite selecting an autosome of similar length, we obtained significantly more SNPs from AUT than from X (**Table S4**). This difference, however, played a subordinate role, as downsampling the autosomal data had little impact on the results (**Supplementary Text 2.4**). Despite this limited impact, we would still advise selecting an autosome that represents genome-wide variation while sharing a similar recombination rate and GC content with the sex chromosome, to partially account for linked selection. To further avoid bias and equalise analytical power, SNP numbers should probably also be matched between datasets.

Sequence coverage is another factor likely to prove decisive, as higher coverage yields more stable estimates of genetic diversity. We worked with low-to-medium coverage data but show that site number and data quality were broadly equivalent at 7x, 10x, and 16x (**Supplementary Text 2.5**). Coverage differences may nonetheless affect diversity estimates, and consistent with the literature, estimates based on genotype likelihoods yielded more stable results. Overall, however, choices of genotyping and filtering strongly affected *Ne^F^/Ne^M^* estimates. For example, a difference in X-chromosomal diversity estimates of only 0.00001 propagated into substantial downstream differences with *Ne^F^/Ne^M^* ratios ranging from 1.085 to 1.411 (GSL, ANGSD θ*^W^*estimates in **Table 6 & 7 in Supplementary Text 2.3**); several combinations of filtering choices and estimators even produced biologically implausible negative estimates.

Our exploration of methodological choices was necessarily opportunistic and far from exhaustive. It does, however, suggest that bioinformatic choices can have a strong impact on estimates of genetic diversity and resulting interpretation, warranting systematic, in-depth evaluation (cf. Shafer et al., 2017).

### Conclusion

We leveraged variation of four genomic compartments with different matri- and patrilineal contributions, including the Y chromosome, to estimate sex bias in reproductive skew and dispersal in two closely related sea lion species. Consistent with expectations from their life histories, both aspects were more pronounced in males, with a stronger signal in the more polygynous species. While our results align with expectations based on the species’ biology and support a possible link between mating system and dispersal, we show that both confounding biological processes and variation introduced during bioinformatic processing can strongly influence the results. A promising way forward is to implement a highly integrative inference framework that jointly models all interfering processes by simulation (Haller & Messer, 2022; Miyagi et al., 2025), ideally repeated across different bioinformatic pipelines to test for robustness. This, however, is not currently possible as it would need a recombination map, direct sex-specific estimates of the mutation rate, and robust genotyping methods, underscoring the continued need for basic genomic research in non-model species.

## Data availability statement

Raw sequence reads and related metadata are deposited in the SRA (Bioproject: PRJNA1073733; reviewer link: https://www.ncbi.nlm.nih.gov/bioproject/1073733). All code can be found on GitHub: https://github.com/EvoBioWolf/PINNIPED_sexbiasedmigration

## Funding statement

This work was funded by LMU Munich (J.B.W.W), the Max Planck Society under the Max-Planck Fellow framework (J.B.W.W), the German Research Foundation grant HO 5122/4-1 to J.I.H., J.B.W.W). C.K was supported by a SNSF Postdoc.Mobility (grant nb: #230483). Computing was performed on the BioHPC hosted at Leibniz Rechenzentrum Munich funded by German Research Foundation grant INST 86/2050-1 FUGG (J.B.W.W.).

## Conflict of interest disclosure

The authors declare no conflict of interest.

## Acknowledgements

We are grateful to Gaby Kumpfmüller for performing DNA extractions and to Tom Gelatt for providing SSL samples. We thank David Aurioles Gamboa for providing samples of the California sea lion used for validation of Y-linked SNPs. We are likewise thankful to Katharina Wittman, Ileana Sáenz-Soto and Saurabh Pophaly who helped explore the feasibility of this project during an early stage.

## Data Accessibility and Benefit-Sharing

Raw sequence reads and related metadata are deposited in the SRA (Bioproject: PRJNA1073733; reviewer link: https://www.ncbi.nlm.nih.gov/bioproject/1073733). The raw sequencing data for MassARRAY sequences are available from the authors upon request. All code can be found on GitHub: https://github.com/EvoBioWolf/PINNIPED_sexbiasedmigration

Benefits Generated: Benefits from this research accrue from the sharing of our data and results on public databases as described above.

## Author contribution

J.B.W.W conceived of the study;

D.A.G, A.B.B, J.W.B, J.I.H, J.B.W.W provided samples;

A.Y, J.I.H, J.B.W.W generated sequencing data;

S.G designed MassARRAY assays and generated MassARRAY genotyping data;

A.Y, C.K, L.B, C.R.P, F.B.C performed bioinformatic analyses;

A.Y, C.K, J.B.W.W wrote the manuscript:

All authors edited and agreed on the final manuscript

## Notes

### Competing Interest Statement

The authors have declared no competing interest.

