## Supplementary figures for "Sex-linked genomic variation reveals reproductive skew and male-biased dispersal in polygynous sea lions"

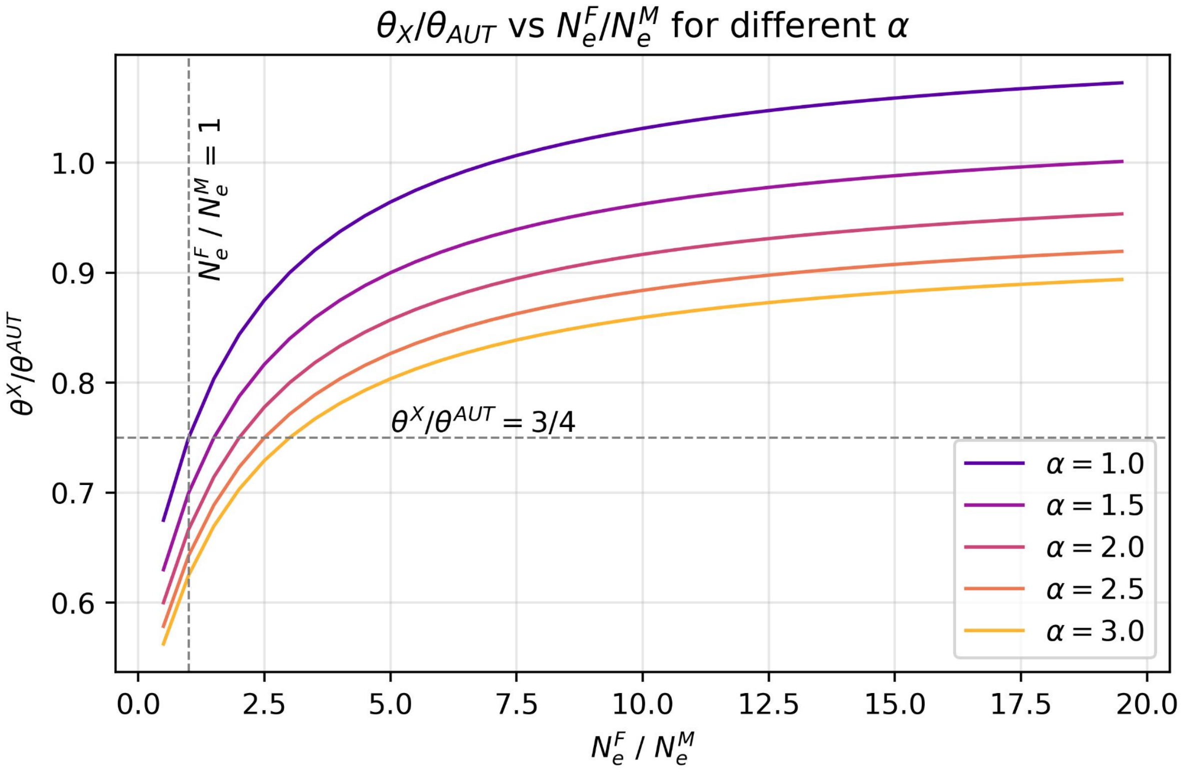


**Figure S1:** Theoretical relationship between effective female-to-male population size ratio ($N_e^F/N_e^M$) and X/AUT diversity with different values for male-biased mutation ($\alpha$ ).

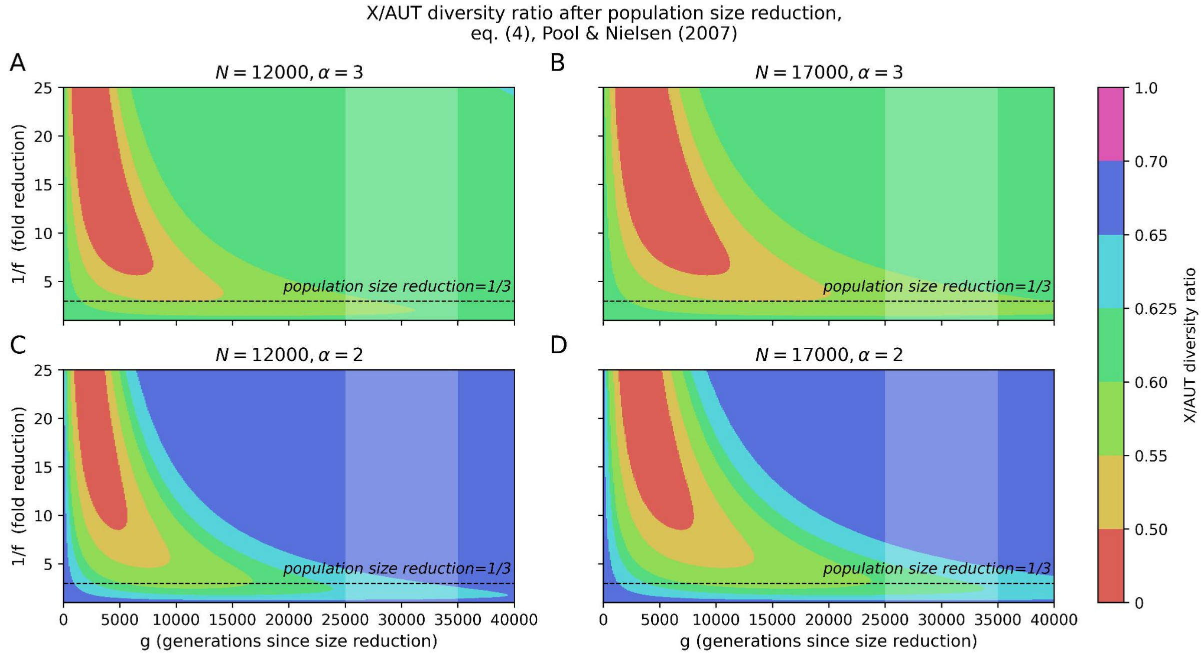


**Figure S2:** Predicted X/A diversity ratio after population size reduction adapted from Pool & Nielsen (2007) with *α=3* (A, B) and *α=2* (C, D) and effective population sizes for GSL ($N_e$*=12000*, A, C) and SSL ($N_e$*=17000*, B, D) estimated as in Peart et al. (2020). The dashed horizontal line indicates a population reduction in up to 30,000 generations after population reduction.

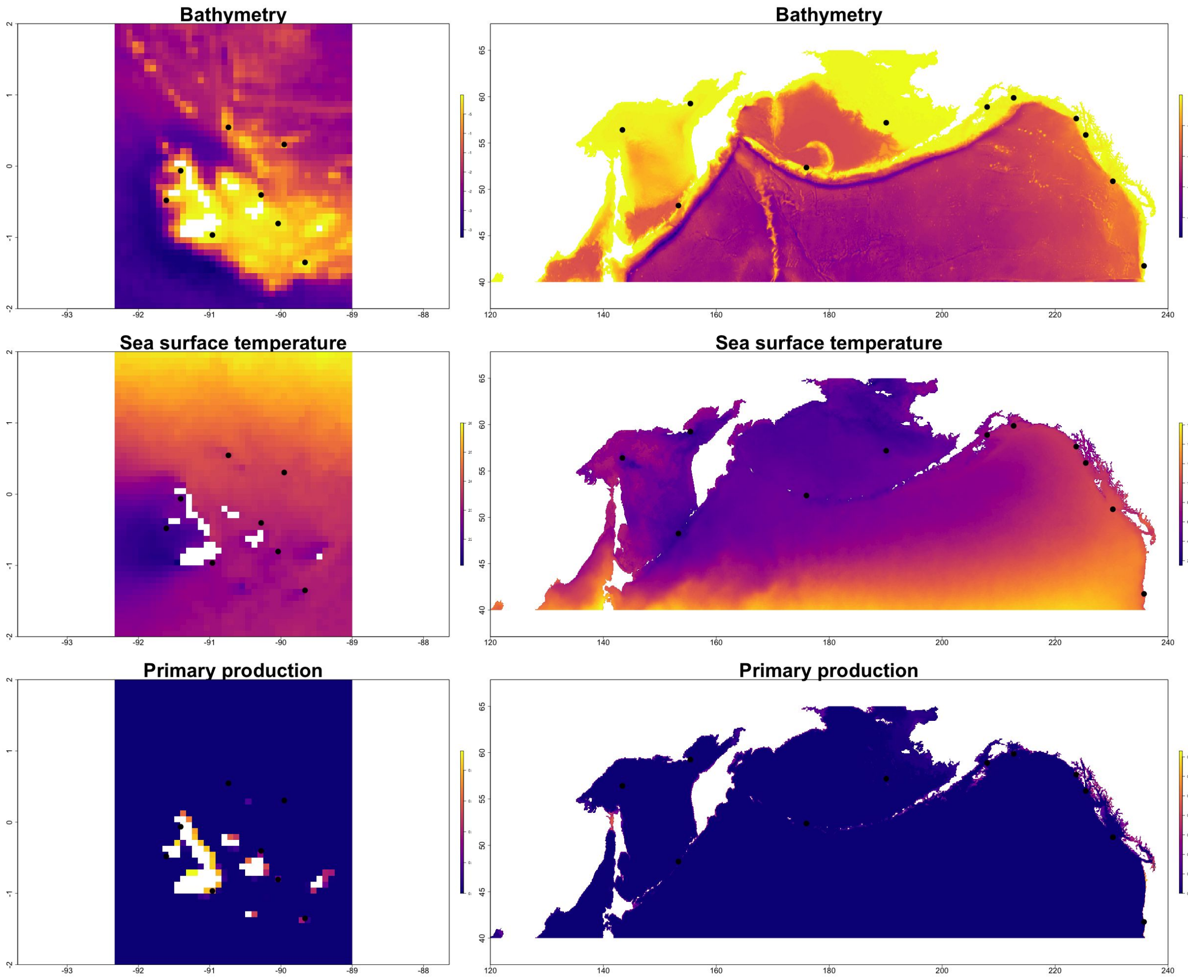


**Figure S3:** Environmental differences across GSL (left) and SSL (right) ranges. Environmental factors taken into account are: sea surface temperature ("MS_biogeo13_sst_mean_5m"), bathymetry ("MS_bathy_5m") and primary productivity ("BO21_ppmean_bdmean"). Black dots represent the sampled rookeries.

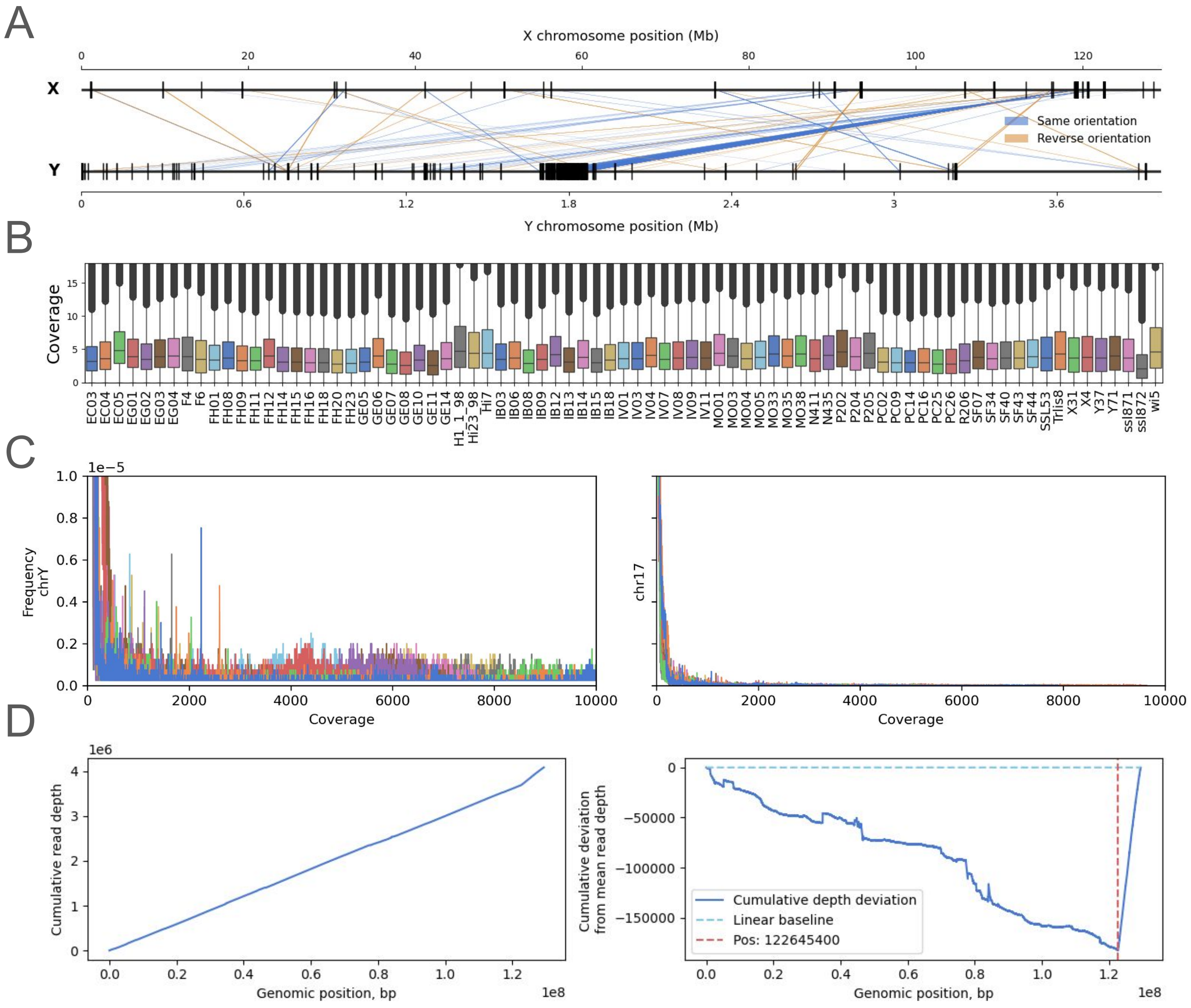


**Figure S4:** Male-specific region of the Y chromosome assembly and X chromosome PAR. **A**: X–Y map of regions with high similarity for the Californian sea lion assembly (repeats masked, identity>90, minimum length=100). **B**: Coverage distribution for Y chromosomal sites with MQ>20 across the Y chromosome for representative samples; **C:** Frequency distribution of sequence coverage on the Y chromosome (left) and chromosome 17 (right) across all samples from panel B. **D**: Identification of the X-chromosome pseudoautosomal boundary in a high-coverage male Galápagos sea lion (FH01), a changepoint, marked with red dashed vertical line, marks the PAR start coordinate.

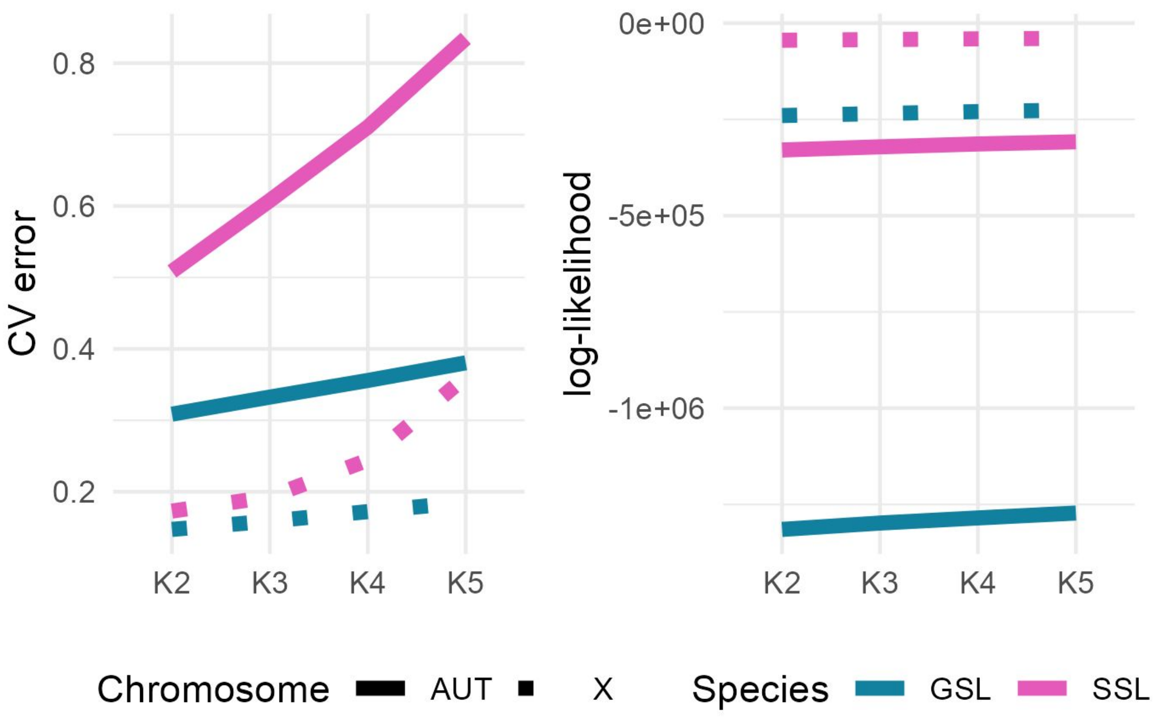


**Figure S5:** ADMIXTURE analysis showing the relationship between the number of ancestral populations (K) and cross-validation (CV) error (left) and log-likelihood (right) for autosomes (solid line) and the X chromosome (dotted line) of both species.

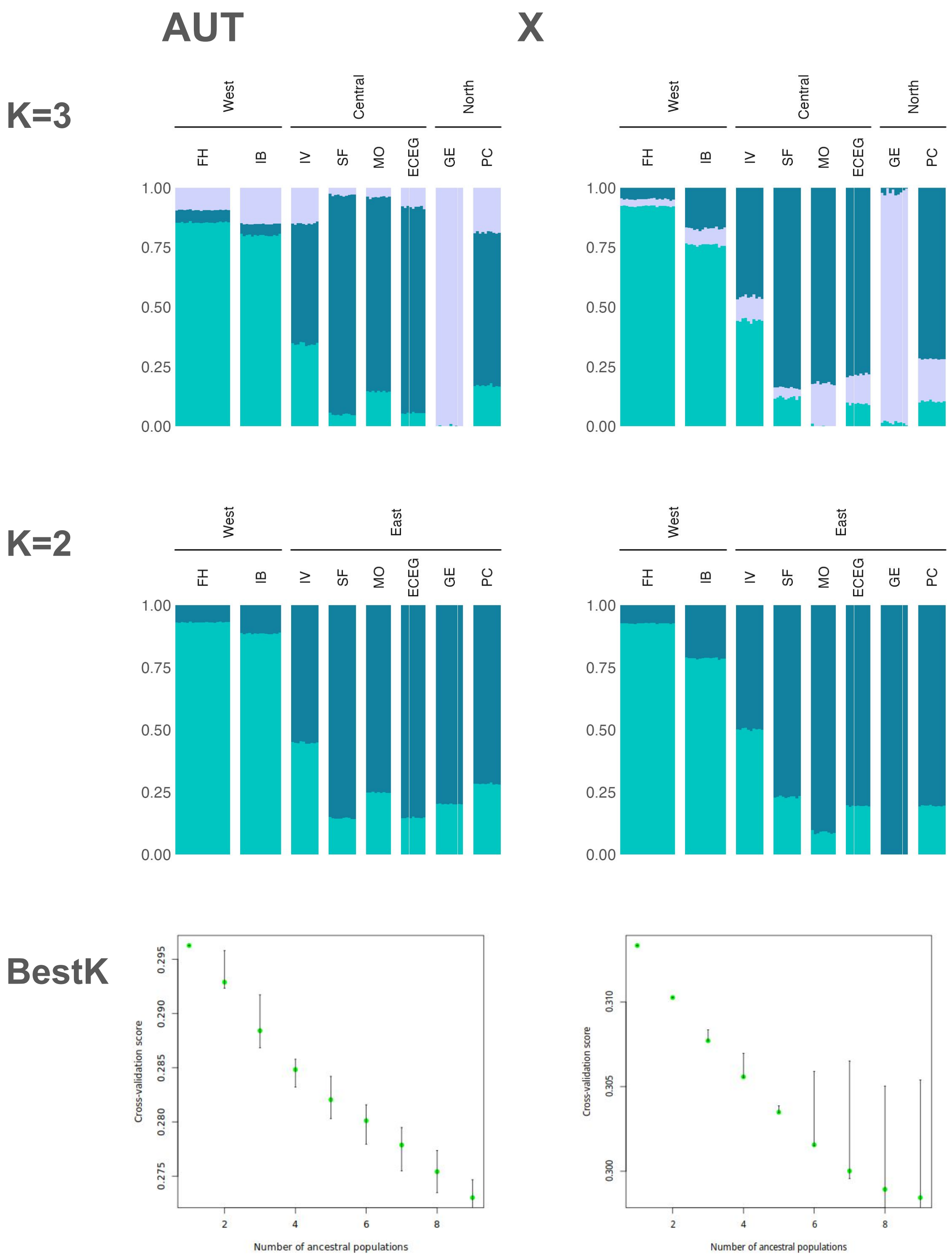


**Figure S6:** Results from spatially explicit structure analyses (TESS) for both the autosome and the X chromosome assuming two or three ancestral populations for the GSL. 'BestK' shows the cross-validation statistics for a varying number of ancestral populations K.

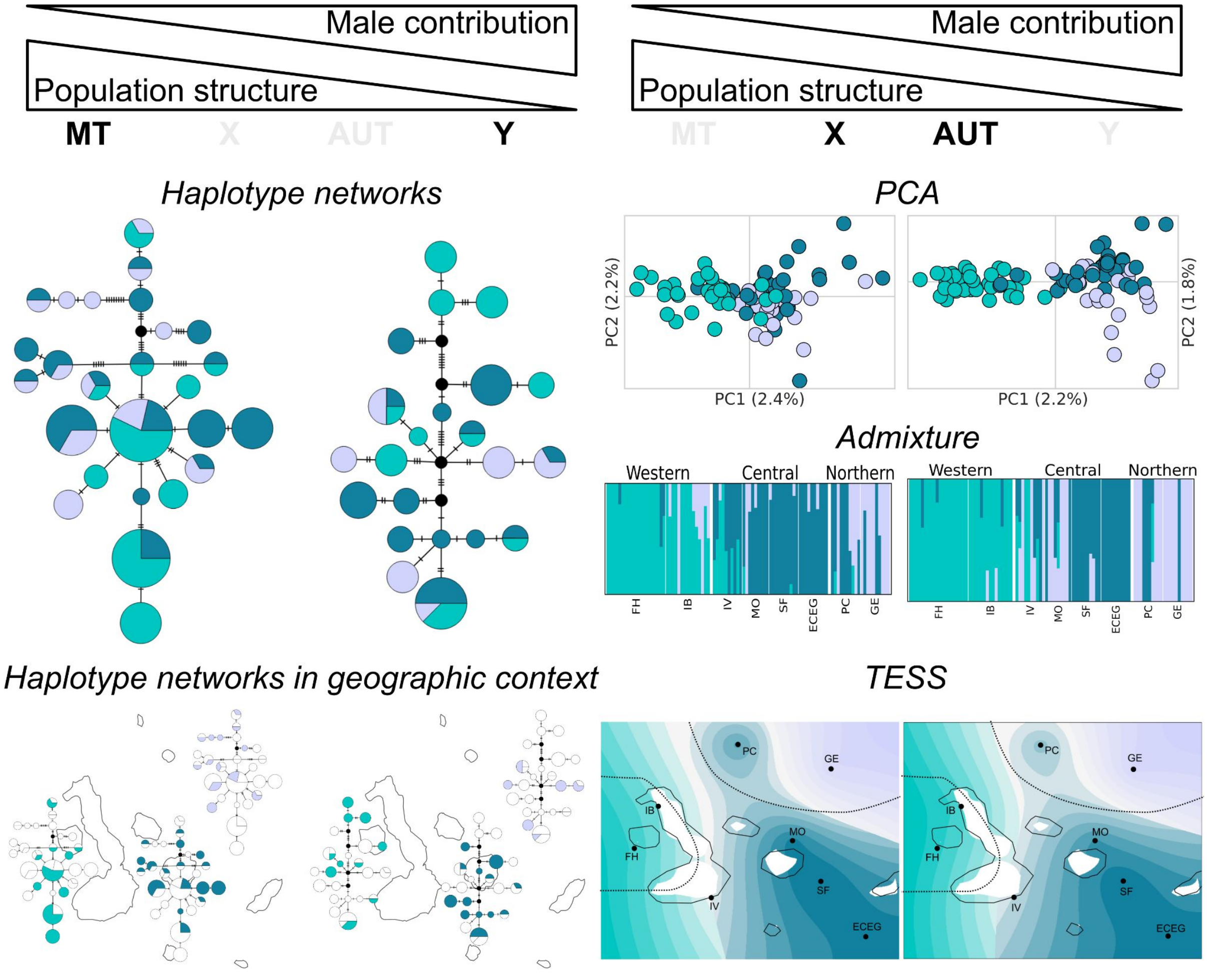


**Figure S7:** Population structure across haploid (MT, Y) (left) and diploid (AUT) or hemizygous (X) (right) genomic compartments in GSL assuming K = 3 ancestral populations. The top row shows expectations regarding the magnitude of population stratification with increasing contribution of male migration. (Left) Haplotype networks are shown for MT and Y where node size represents number of individuals. Haplotype network are also overlaid on the map highlighting haplotypes occurring in each of the three stocks; (Right) PCA for both AUT and X with circles coloured by stocks. Below ancestry proportions from ADMIXTURE analyses, and their geogrhapic interpolation from TESS. Abbreviation refer to rookeries, the dashed line separates stocks. Colours represent stocks as defined in the literature for haplotypes and PCA and as assigned by the analysis in TESS and admixture.

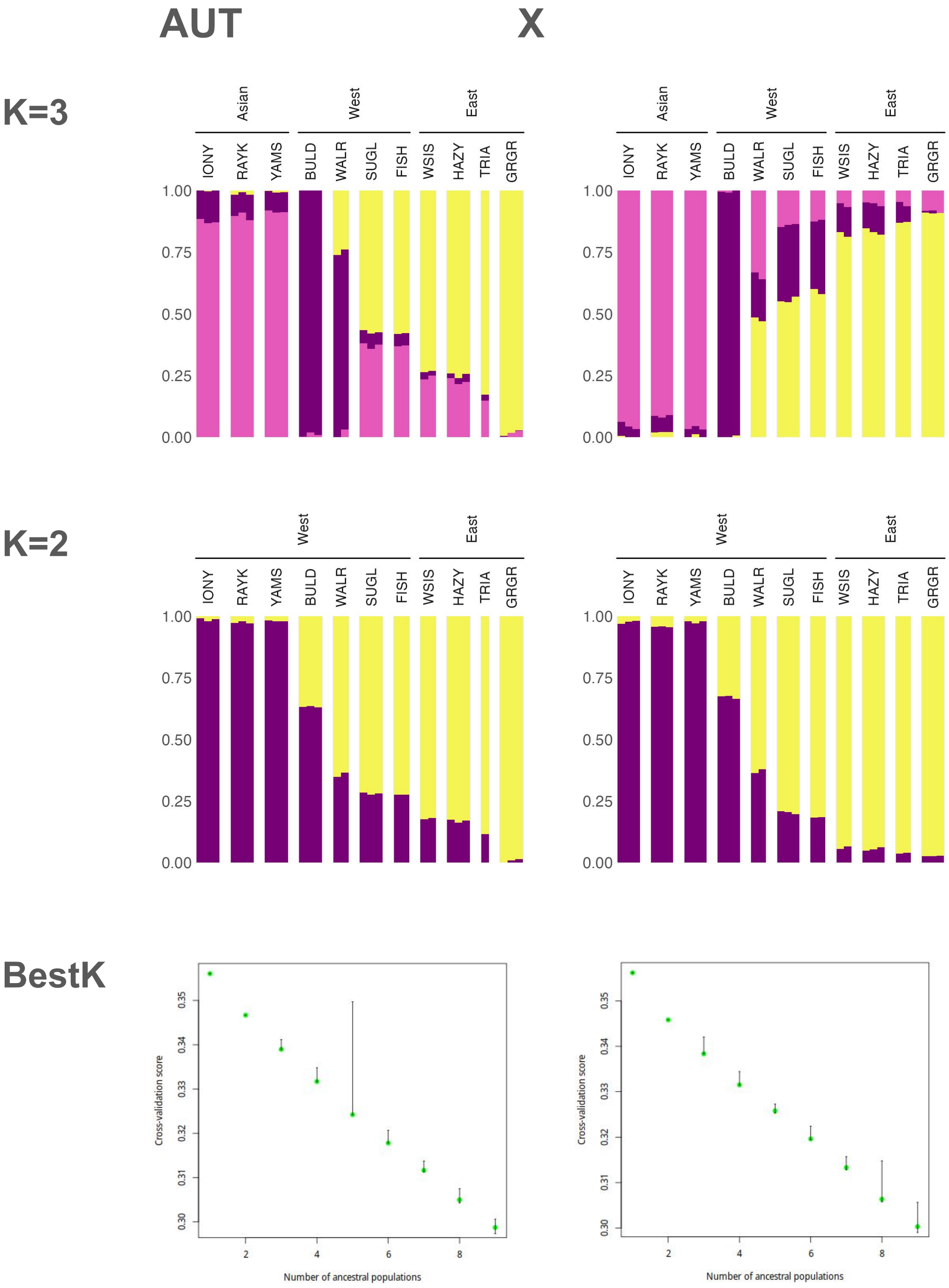


**Figure S8:** Results from spatially explicit structure analyses (TESS) for both the autosome and the X chromosome assuming two or three ancestral populations for the SSL. 'BestK' shows the cross-validation statistics for a varying number of ancestral populations K.

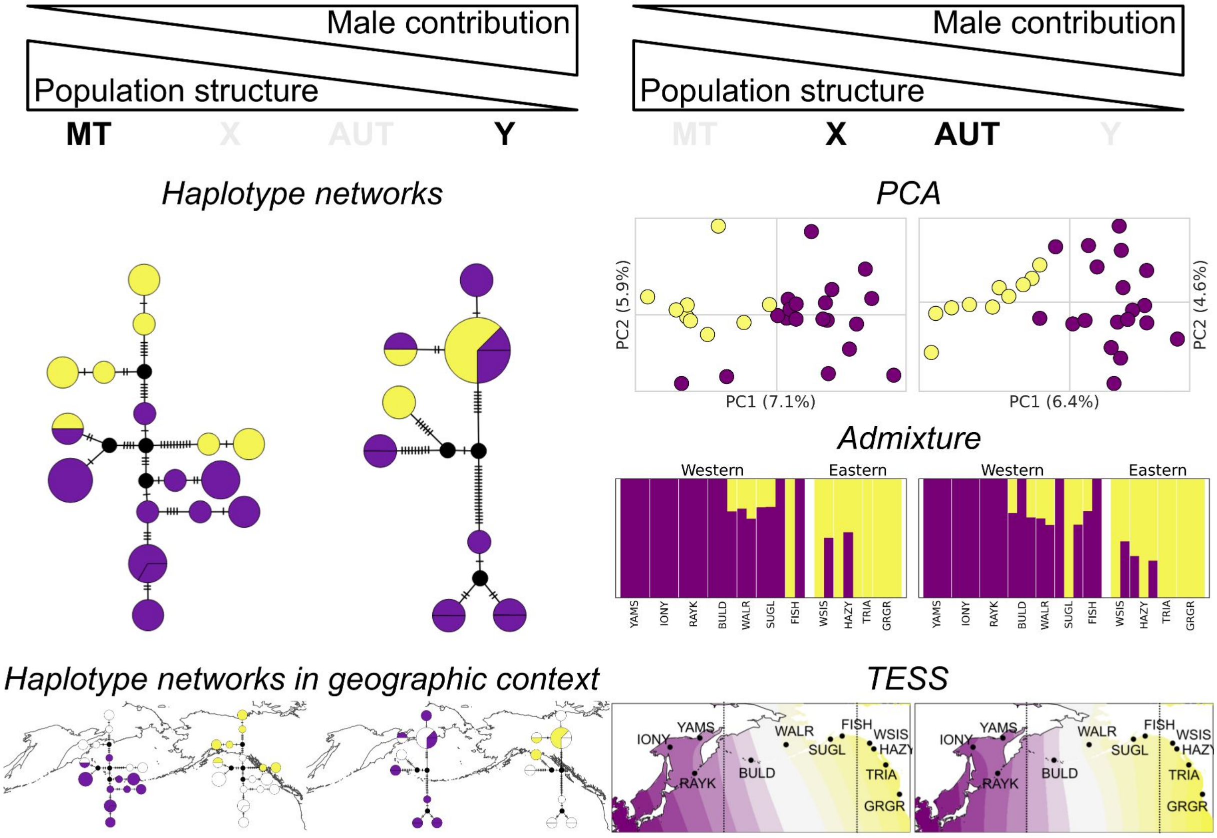


**Figure S9:** Population structure across haploid (MT, Y) (left) and diploid (AUT) or hemizygous (X) (right) genomic compartments in SSL assuming K = 2 ancestral populations. The top row shows expectations regarding the magnitude of population stratification with increasing contribution of male migration. (Left) Haplotype networks are shown for MT and Y where node size represents number of individuals. Haplotype network are also overlaid on the map highlighting haplotypes occurring in each of the three stocks; (Right) PCA for both AUT and X with circles coloured by stocks. Below ancestry proportions from ADMIXTURE analyses, and their geogrhapic interpolation from TESS. Abbreviation refer to rookeries, the dashed line separates stocks. Colours represent stocks as defined in the literature for haplotypes and PCA and as assigned by the analysis in TESS and admixture.

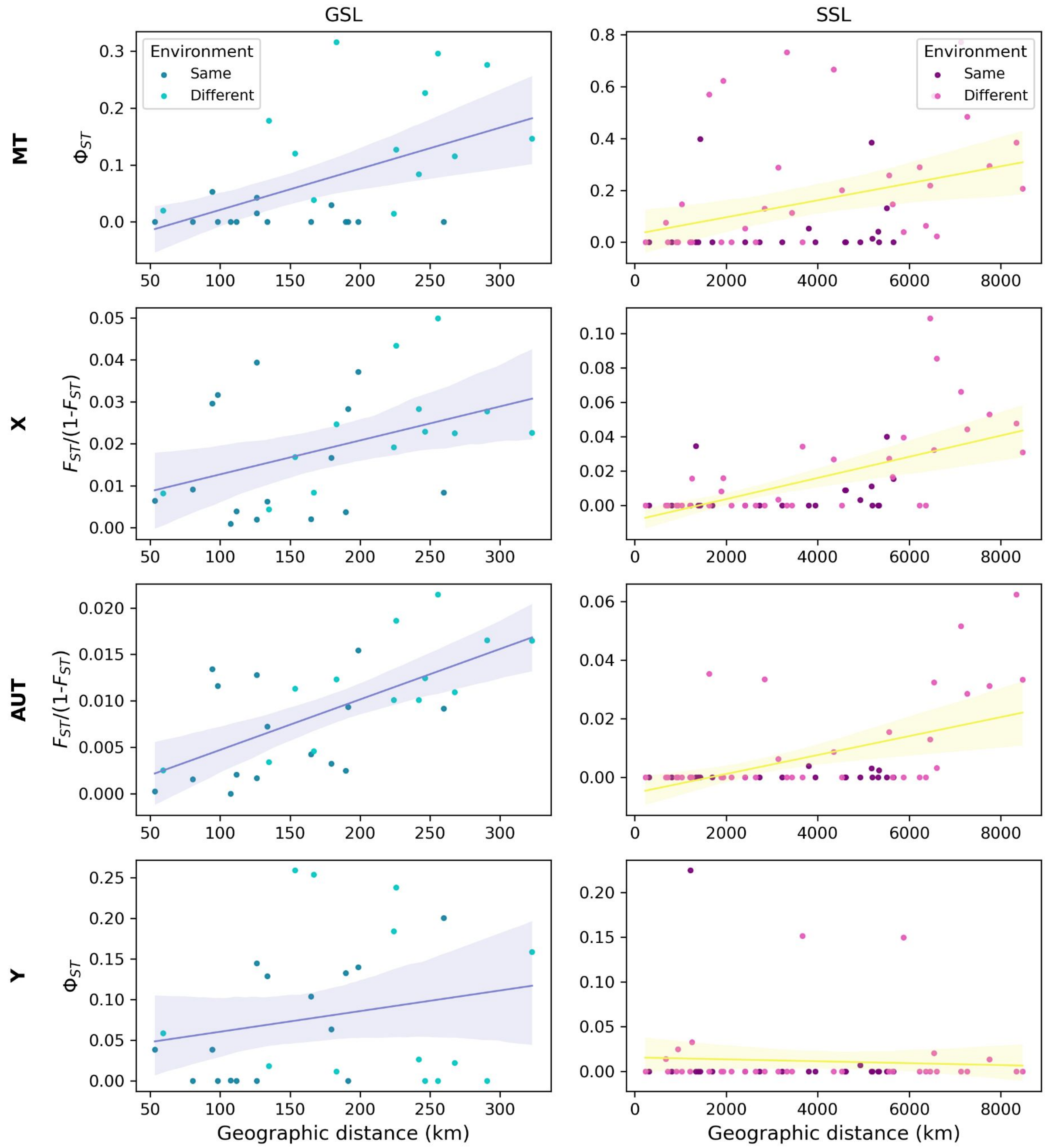


**Figure S10:** Isolation by distance (IBD) in GSL (left) and SSL (right) color coding population comparison within the same or different habitat. Hudson's Fst with Slatkin's linearisation vs geographical distance for each compartment. Linear regression and 95 CI, points are coloured as whether the compared rookeries are in the same or different environment (west vs center/north for GSL & east vs asia/west for SSL).

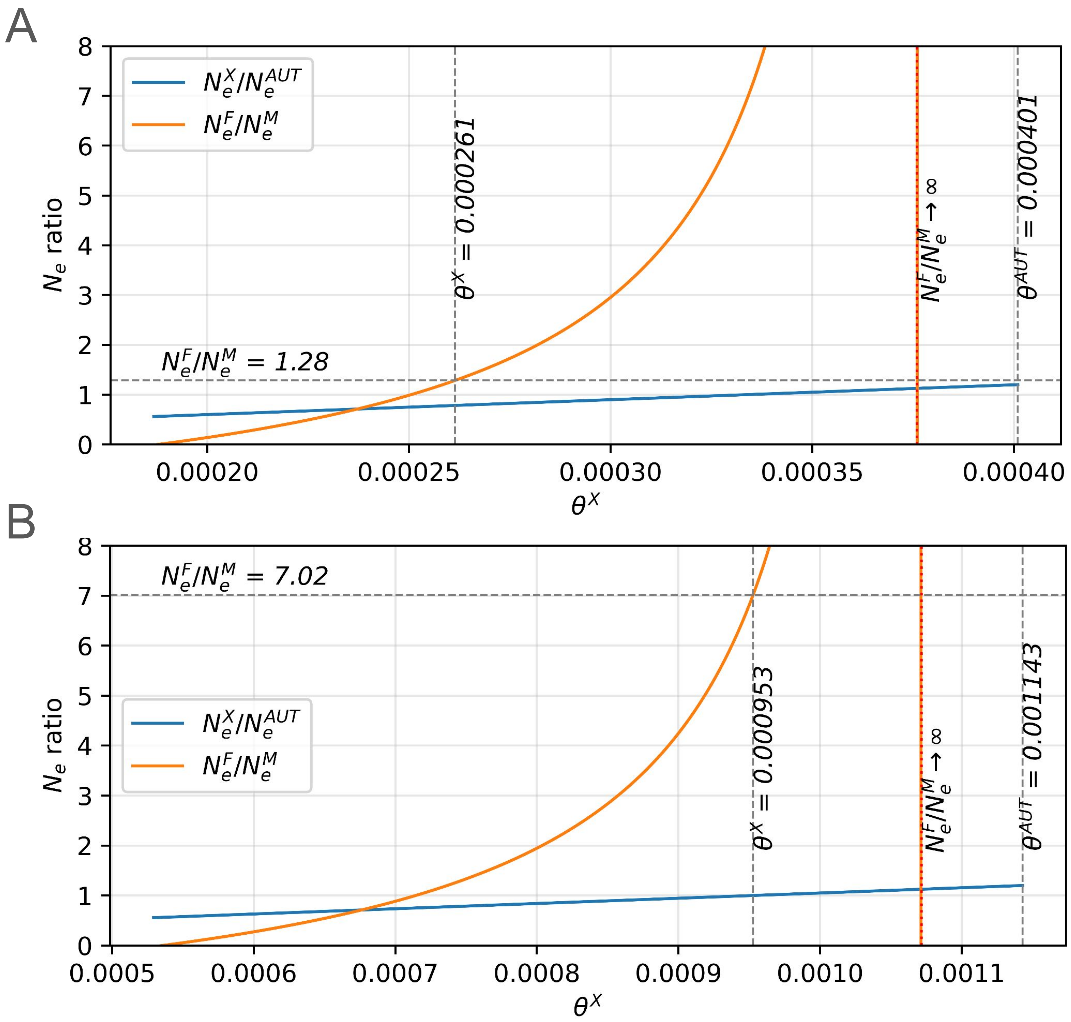


**Figure S11:** Expected ratios of effective population sizes between males and females ($N_e^F/N_e^M$ orange solid line) and between the X chromosome and autosomes ($N_e^X/N_e^{AUT}$ blue solid line) are plotted as a function of X-linked diversity $\theta^X$ for (**A**) the GSL and (**B**) the SSL. The red dashed vertical line marks the asymptote of $N_e^F/N_e^M$. Grey dashed vertical lines indicate the observed X-linked and AUT diversity ($\theta^X$ and $\theta^{AUT}$, respectively). The grey dashed horizontal line indicates the estimated effective female-to-male population size ratio ($N_e^F/N_e^M$).
