## supplementary text for "Sex-linked genomic variation reveals reproductive skew and male-biased dispersal in polygynous sea lions"

### Exploiting genomic variation in sex-linked inheritance to assess sex-biased dispersal in sea lions - Supplementary text

A. Yakupova & C. Kessler

November 2, 2026

### 1 MassARRAY assay development and genotyping.

MassARRAY assays were developed before the current VGP genome was available. For that approach, the assembly was generated using CLC Genomics Workbench v9.5.3 (CLCbio, Qiagen) with default parameters. To identify Y chromosome scaffolds, we applied the average depth ratio (AD-ratio) method (Bidon et al., 2014). This method distinguishes autosomal, X, and Y chromosome scaffolds based on the relative female-male read depth. We sequenced short-read libraries from a female (15.9x) and a male (14.2x) California sea lions (CSL) (Table S2, MA_SI), and mapped them to the male genome assembly using BWA MEM v0.7.8 (H. Li, 2011). Then, we used Picard v2.0.1 (BroadInstitute, 2019) to add read groups, sort and remove duplicate reads, and merge the BAM files from separate sequencing runs. Reads were realigned around indels with GATK v3.7 (McKenna et al., 2010; Van Der Auwera et al., 2013), and reads with MQ $< 20$ and scaffolds shorter than 1kb or no mapped reads were excluded from further analyses. For the average read depth calculation, we also removed positions with missing data and those with coverage $\leq$ 50x. We used samtools depth v1.3 (Li and Durbin, 2009) to compute the depth ratio, which we normalised for differences in sequencing coverage, and aligned all obtained putative Y chromosome scaffolds (AD-ratio $\leq 0.2$) to the female genome assembly. Putative Y chromosome scaffolds with 95% sequence identity to the female genome were excluded from the assembly.

To validate Y-chromosomal genotype calls and reduce the high rate of missing data resulting from low Y-scaffold coverage, we developed a genotyping assay. To identify candidate segregating variants on the Y, we first aligned data from 59 male and five female CSL and GSL samples (Table S2, MA_YD) as described above, considering only primary alignments with MQ $> 20$, and using the female genome assembly (V1) supplemented with the validated Y-linked scaffolds ($n = 648$) as reference.

We restricted our variant selection to single-copy genes within the male-specific Y chromosome region (scMSY) (Wallner et al., 2017) to prevent false positives in sequences derived from ampliconic Y regions. To do this, we manually extracted exons of seven known dog scMSY genes (USP9Y, UBE1Y, DDX3Y, ZFY, KDM5D, EIF1AY, UTY; Genbank accession: KP081776) using Geneious v. 11.0.5 (Kearse et al., 2012), to which we aligned the pool of putative Y scaffolds with CLC Main Workbench 7.7.3 (CLCbio, Qiagen) and default settings. We manually inspected the alignments and considered scaffolds homologous if they lacked stop codons and excessive mismatches. Then, we used samtools depth to calculate average scaffold coverage for all homologous scaffolds ($n = 35$), which we used to calculate the mean normalised average coverage (MNAC) across all male samples. We retained scaffolds with an MNAC $\leq 1.5$ for the downstream analysis.

Lastly, we performed variant calling per sample using GATK HaplotypeCaller (v3.7) in diploid mode, followed by joint genotyping with GenotypeGVCFs. Variants were excluded with BCFtools v1.9 (Li et al., 2009), BEDOPS v2.4.3 (Neph et al., 2012), bedtools v2.27.1 (Quinlan and Hall, 2010) and VCFtools v0.1.15 (Danecek et al., 2011) based on the following criteria (Wallner et al., 2017): (1) presence in female samples; (2) heterozygous calls in males; (3) fixed alternative alleles; (4) coverage $< 2$ reads per individual; (5) quality genotypes $< 70$; (6) presence in $< 50\%$ of males; (7) total coverage $> 210$ across all samples; and (8) variants within 100 bp of another variant. We also removed indels and masked non-target variants and repetitive regions identified with RepeatMasker v4.0.7 (Smit et al., 2015), resulting in a final dataset of 72 high-confidence Y-linked SNPs.

For this set of 72 Y-linked candidate SNPs, we designed genotyping assays consisting of forward, reverse, and extension primers using the MassARRAY® System (Agena Bioscience®; Ellis and Ong, 2017). Primers were compared cross-validated to the female sea lion genome (V2) using BLAST v2.7.1+ (Camacho et al., 2009) to assess potential non-specific amplification. We genotyped 62 sea lions (Table S2, MA), including five female samples and one water blank as controls. To assess reproducibility, ten male samples were genotyped in triplicate and nine in duplicate. Assay design and genotyping were performed at the Institute of Clinical Molecular Biology of the Christian-Albrechts-University Kiel.

Genotypes were manually curated based on a visual assessment of signal output strength. We excluded assays that showed poor or non-specific amplification in the blank control and in female samples. Mismatches between assay-derived and bioinformatically inferred genotypes, as well as between technical replicates, were attributed to laboratory contamination and were excluded along with samples with excessive missingness. Finally, to assess the reliability of the genotyped Y-markers, we used BLAST to compare the sequences used for assay design with the male sea lion genome, which was not available at the time of assay design. The best BLAST hit was consistently found on the Y chromosome. Several sequences showed the best matches with different regions of the Y chromosome and were excluded. For GSL, our final set yielded 27 high-confidence Y-linked species-specific SNPs, genotyped in 94 male GSL individuals.

#### 2 Impact of data filtering

##### 2.1 Effect of autosomes reduction to a single representative

Throughout this study, chromosome CM019810.2 from the CSL genome assembly, corresponding to chromosome 9, was used as a representative autosome for comparisons with chromosome X. To evaluate whether this chromosome is representative of genome-wide autosomal patterns, we compared its population structure and levels of genetic diversity to those observed in the remaining autosomes (Figure 1, 2).

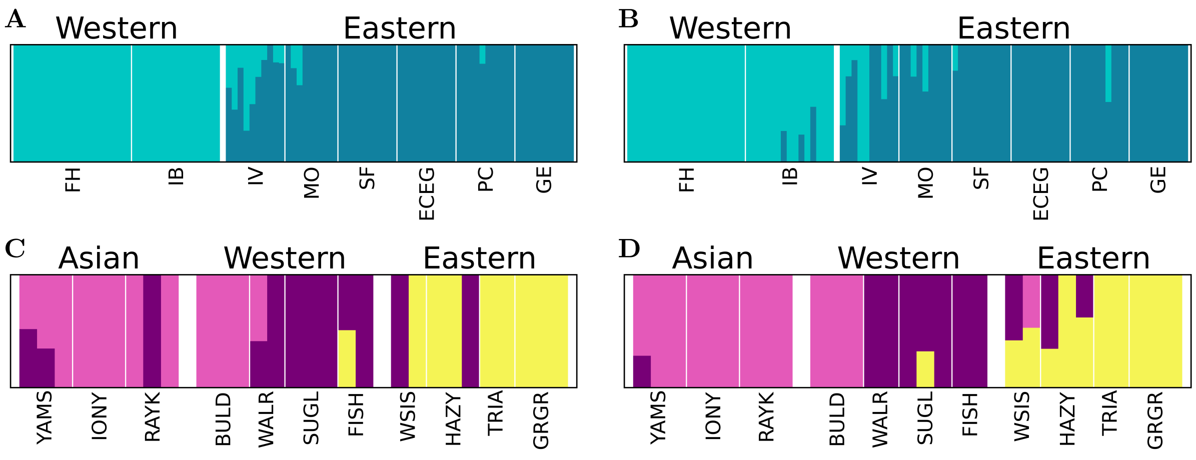

Figure 1: ADMIXTURE proportions comparisons for GSL (**A**, **B**) and SSL (**C**, **D**) based on the data from all autosomes combined (**A**, **C**) and a single autosomal representative CM019810.2 (**B**, **D**).

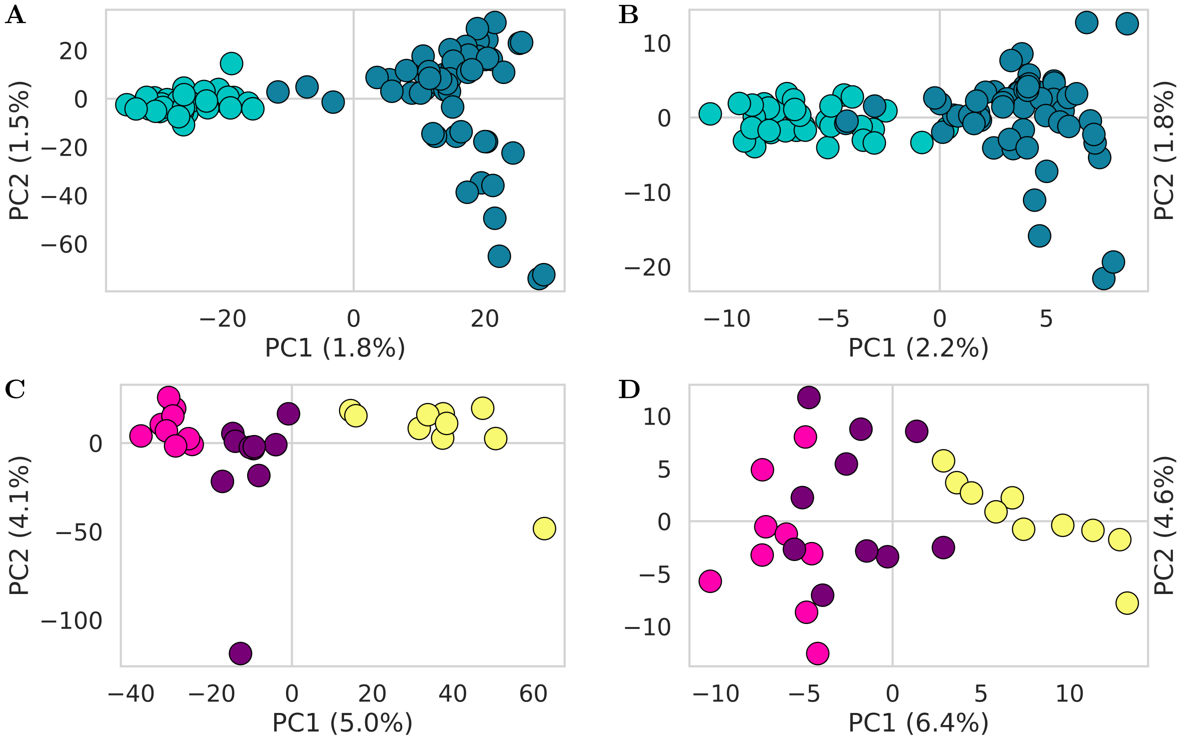

Figure 2: PCA comparisons for GSL (**A**, **B**) and SSL (**C**, **D**) based on the data from all autosomes combined (**A**, **C**) and a single autosomal representative CM019810.2 (**B**, **D**).

To evaluate whether chromosome CM019810.2 was significantly different in genetic diversity, we compared its values of $\theta_W$ and $\pi$ with those of the remaining autosomes (Table 1). In both GSL and SSL, CM019810.2 fell well within the distribution of autosomal diversity estimates and was not identified as an outlier by either Z-score or interquartile range (IQR) criteria (Table 2).

During this analysis, we noticed that chromosome CM019806.2, identified as chromosome 5 in the CSL genome assembly annotation, showed elevated diversity relative to all other autosomes. Both $\theta_W$ and $\pi$ were 4-7-fold higher than the genomic background in both species, resulting in extreme Z-scores and was also classified as an outlier under the IQR criterion (Table 3). We repeated the comparison after excluding CM019806.2, because it strongly inflated the overall variance. The resulting distributions were considerably narrower, and CM019810.2 remained close to the autosomal mean and with $|Z| < 0.6$ for all comparisons (Table 4), further supporting that it is not an outlier.

All comparisons performed show that CM019810.2 is not atypical in terms of diversity or differentiation and could not bias downstream analyses based on a single autosomal chromosome.

Table 1: Summary statistics, calculated with ANGSD ($\pi$ and $\theta_W$) for all autosomes separately and combined for GSL and SSL

| chr. | $\theta_W$ (GSL) | $\pi$ (GSL) | $\theta_W$ (SSL) | $\pi$ (SSL) |
|---|---|---|---|---|
| CM019802.2 | 0.000363 | 0.000292 | 0.000981 | 0.000681 |
| CM019803.2 | 0.000368 | 0.000310 | 0.001007 | 0.000717 |
| CM019804.2 | 0.000378 | 0.000321 | 0.001014 | 0.000714 |
| CM019805.2 | 0.000370 | 0.000285 | 0.000987 | 0.000677 |
| CM019806.2 | 0.001630 | 0.002400 | 0.004901 | 0.006224 |
| CM019807.2 | 0.000364 | 0.000292 | 0.000997 | 0.000698 |
| CM019808.2 | 0.000394 | 0.000354 | 0.001016 | 0.000730 |
| CM019809.2 | 0.000386 | 0.000304 | 0.001003 | 0.000694 |
| CM019810.2 | 0.000410 | 0.000372 | 0.001028 | 0.000737 |
| CM019811.2 | 0.000449 | 0.000314 | 0.001141 | 0.000833 |
| CM019812.2 | 0.000415 | 0.000363 | 0.001043 | 0.000735 |
| CM019813.2 | 0.000380 | 0.000328 | 0.001004 | 0.000705 |
| CM019814.2 | 0.000408 | 0.000326 | 0.000998 | 0.000692 |
| CM019815.2 | 0.000535 | 0.000468 | 0.001217 | 0.000952 |
| CM019816.2 | 0.000438 | 0.000370 | 0.001118 | 0.000811 |
| CM019817.2 | 0.000484 | 0.000415 | 0.001198 | 0.000928 |
| CM019818.2 | 0.000481 | 0.000408 | 0.001171 | 0.000887 |
| AUT | 0.000485 | 0.000466 | 0.001284 | 0.001083 |

Table 2: Chromosome CM019810.2 exhibits diversity levels comparable to the autosomal background in GSL and SSL.

| chr. | $\theta_W$ (GSL) | $\pi$ (GSL) | $\theta_W$ (SSL) | $\pi$ (SSL) |
|---|---|---|---|---|
| CM019810.2 | 0.000410 | 0.000372 | 0.001028 | 0.000737 |
| AUT mean | 0.000490 | 0.000472 | 0.001300 | 0.001105 |
| ± std | ± 0.000308 | ± 0.000517 | ± 0.000964 | ± 0.001368 |
| Z-score | -0.26 | -0.19 | -0.28 | -0.27 |

Table 3: Outlier analysis of chromosome CM019806.2 based on chromosome-wide estimates of $\theta_W$ and $\pi$ in GSL and SSL.

| chr. | $\theta_W$ (GSL) | $\pi$ (GSL) | $\theta_W$ (SSL) | $\pi$ (SSL) |
|---|---|---|---|---|
| CM019806.2 | 0.001630 | 0.002400 | 0.004901 | 0.006224 |
| AUT mean | 0.000414 | 0.000345 | 0.001058 | 0.000762 |
| ± std | ± 0.000051 | ± 0.000052 | ± 0.000082 | ± 0.000091 |
| Z-score | 23.93 | 39.72 | 47.12 | 60.00 |

Table 4: Chromosome CM019810.2 exhibits diversity levels comparable to the autosomal background in GSL and SSL, with lower variance after removing the outlier (CM019806.2).

| chr. | $\theta_W$ (GSL) | $\pi$ (GSL) | $\theta_W$ (SSL) | $\pi$ (SSL) |
|---|---|---|---|---|
| CM019810.2 | 0.000410 | 0.000372 | 0.001028 | 0.000737 |
| AUT mean | 0.000414 | 0.000343 | 0.001060 | 0.000764 |
| ± std | ± 0.000053 | ± 0.000053 | ± 0.000084 | ± 0.000094 |
| Z-score | -0.08 | 0.54 | -0.38 | -0.29 |

#### 2.2 Effect of pseudohaploidisation

As the spatially informed population structure in TESS3R does not allow for mixed-ploidy data, we decided to pseudo-haploidise the X dataset (see main text). To assess whether this approach was consistent with the original variant-calling procedure, we generated admixture proportions and PCA plots from the pseudo-haploid datasets and compared them with those obtained from the original diploid calls.

Those comparisons do not point to number of sites being the source of the lower population structure in X and show similar structure between both baselines and pseudo-haploidised datasets (Figures 3 : 6).

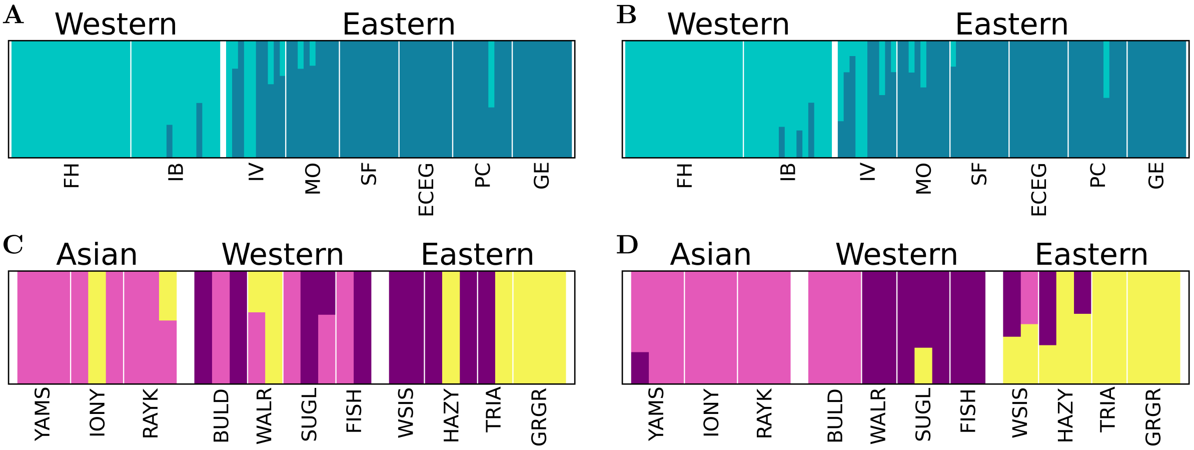

Figure 3: ADMIXTURE proportions comparisons in CM019810.2 for GSL (**A**, **B**) and SSL (**C**, **D**) based on pseudohaplodised (**A**, **C**) and baseline data (**B**, **D**).

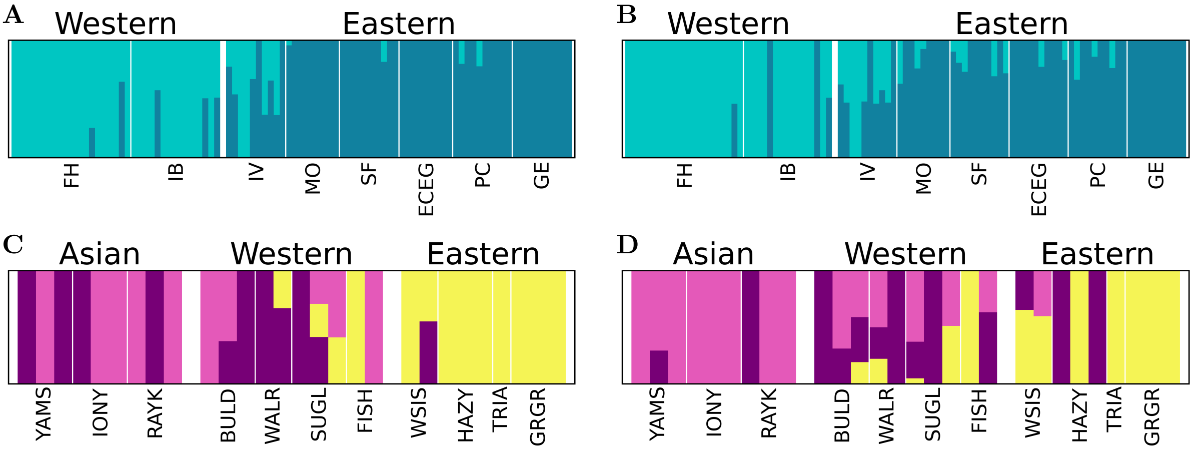

Figure 4: ADMIXTURE proportions comparisons in CM019819.2 for GSL (**A**, **B**) and SSL (**C**, **D**) based on pseudohaplodised (**A**, **C**) and baseline data (**B**, **D**).

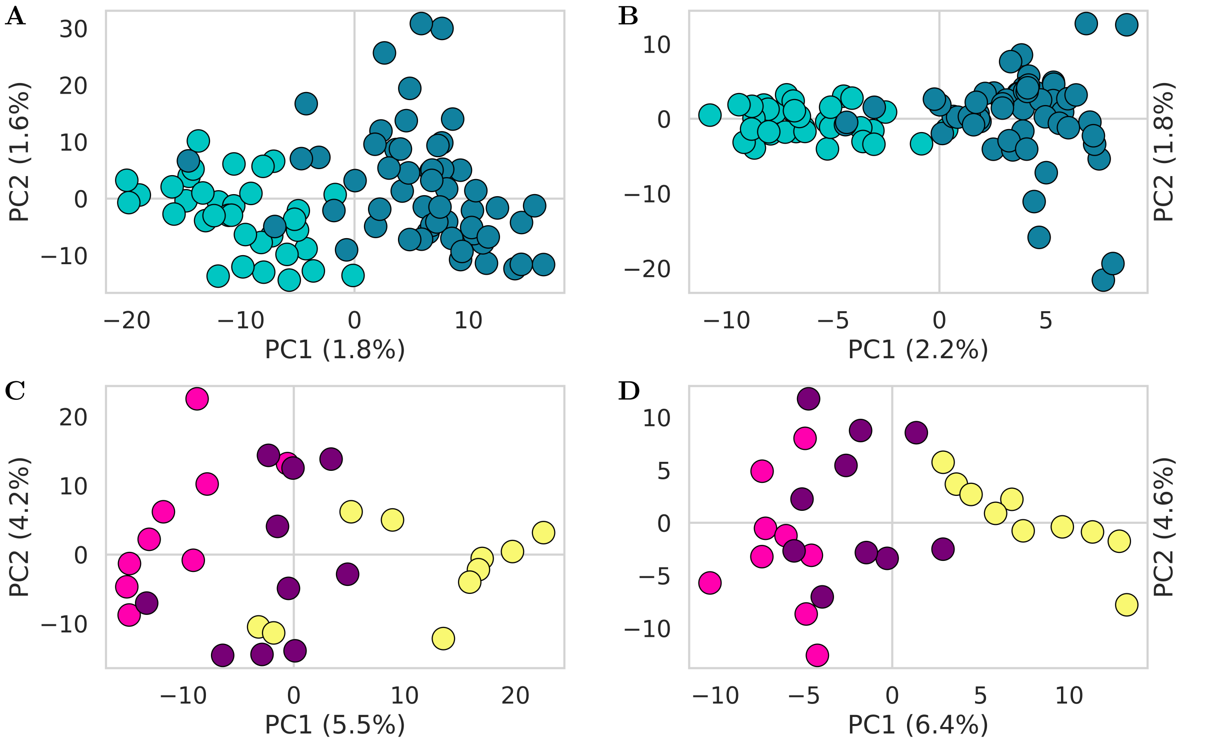

Figure 5: PCA comparisons in CM019810.2 for GSL (**A**, **B**) and SSL (**C**, **D**) based on pseudohaplodised (**A**, **C**) and baseline data (**B**, **D**).

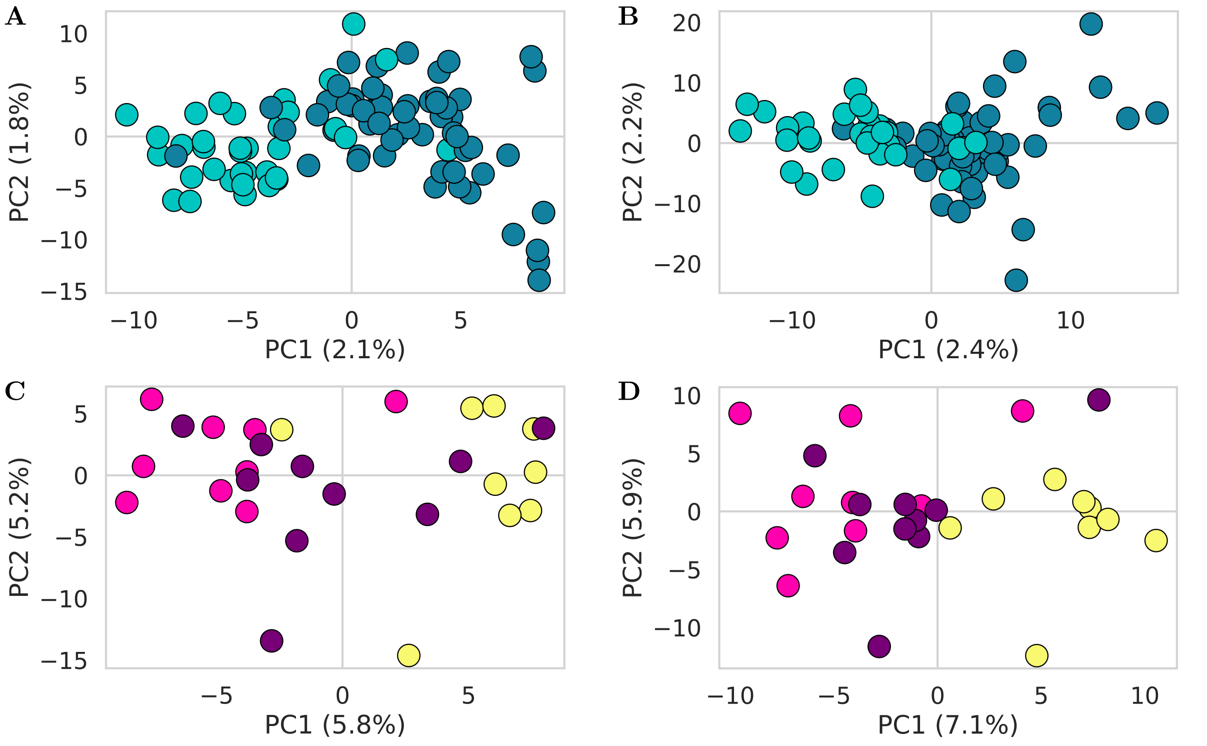

Figure 6: PCA comparisons in CM019819.2 for GSL (**A**, **B**) and SSL (**C**, **D**) based on pseudohaplodised (**A**, **C**) and baseline data (**B**, **D**).

#### 2.3 Effect of filtering on Watterson's theta ($\theta_W$) and genetic diversity ($\pi$) estimations

As part of our initial data exploration, we estimated $\pi$ using Pixy (Korunes and Samuk, 2021). The analyses was based on a dataset that included invariant sites and was generated using the same variant calling and filtering pipeline described in the main text, except for the *-v* flag in *bcftools call.* This dataset was therefore built with several constraints that could compromise the accuracy of $\pi$ estimates, primarily by disproportionately affecting the number of invariant sites (Brault et al., 2026). Although the resulting $\pi$ estimates appeared plausible, the ratios between them deviated substantially from the theoretical expectations (Figure S1), making the corresponding effective population size ($N_e$) ratios unreliable (Table 5).

Table 5: Pixy estimates based on variant and invariant sites, called on males and females, filtered for strand-bias, depth, mapping quality (40) and missing data (10%), estimated across all individuals.

| | GSL | SSL |
|---|---|---|
| $\pi^{AUT}$ | 0.000297 | 0.000484 |
| $\pi^{X}$ | 0.000122 | 0.000199 |
| $\pi^{X}$ / $\pi^{AUT}$ | 0.41116 | 0.41081 |
| $\boldsymbol{N_e^F / N_e^M}$ | **-0.21884** | **-0.22002** |

These results prompted us to reconsider the methodology. We decided to estimate $\theta_W$ alongside $\pi$ and to calculate both summary statistics using ANGSD (Korneliussen et al., 2014). Indeed, both statistics are particularly sensitive to data filtering (Brault et al., 2026). However, ANGSD accounts for genotype uncertainty through genotype likelihoods. Here, given the specific filtering requirements, we first called all sites in males and females using bcftools, and then supplied the resulting site positions to ANGSD using the *-sites* option.

Two additional factors that impacted our estimates were the strand bias filter and the inclusion of males in the X chromosome analysis. The stand bias filter retained only positions at which $< 25\%$ of genotypes exhibited strand bias in MT and $< 50\%$ in AUT, X and Y. Including males in the calculation of summary statistics resulted in a higher proportion of missing data in X than in AUT, potentially introducing disproportionate bias into the X chromosome $\pi$ estimates and, consequently, the inferred $N_e$ ratios.

We therefore removed the strand bias filter, and computed diversity on females only (for both X and AUT), and obtained the following results:

Table 6: Estimates based on variant and invariant sites, called on males and females, filtered for maximum depth, mapping quality (40) and missing data (10%), estimated in females only.

| | Estimated in Pixy | | Estimated in ANGSD | |
|---|---|---|---|---|
| | GSL | SSL | GSL | SSL |
| $\pi^{AUT}$ | 0.00034 | 0.00051 | 0.00037 | 0.00084 |
| $\pi^{X}$ | 0.00014 | 0.00027 | 0.00018 | 0.00061 |
| $\pi^{X}$ / $\pi^{AUT}$ | 0.42884 | 0.52696 | 0.47293 | 0.72480 |
| $\boldsymbol{N_e^F / N_e^M}$ | **-0.15691** | **0.28357** | **0.01799** | **2.40760** |
| $\theta_w^{\mathrm{AUT}}$ | 0.00037 | 0.00216 | 0.00041 | 0.00127 |
| $\theta_w^{X}$ | 0.00023 | 0.00279 | 0.00026 | 0.00108 |
| $\theta_w^{X} / \theta_w^{\mathrm{AUT}}$ | 0.60283 | 1.28861 | 0.63365 | 0.85095 |
| $\boldsymbol{N_e^F / N_e^M}$ | **0.80127** | **-4.67008** | **1.08542** | **8.83231** |

Here, Pixy estimates continued to yield implausible $N_e$ ratios. Although ANGSD estimates did not produce negative ratios, the GSL ratio based on $\pi$ suggests an extreme female bias (about 100 females per male), whereas the ratio based on $\theta_W$ indicated an approximately equal sex ratio. Neither result is consistent with GSL biology. For SSL, the ratios obtained using ANGSD were more biologically plausible.

As described above, all sites were initially called using bcftools and then the obtained positions were supplied to ANGSD, meaning that the ANGSD estimates were still influenced by the regular variant calling method. We also noticed that including males in the genotype calling resulted in a greater proportion of sites being removed from X than from AUT. Therefore, for analysis in Pixy, we re-ran the genotype calling using only female individuals. The simplified filtering strategy allowed us to apply filters directly in ANGSD, eliminating the need for prior variant calling and filtering. Here, we applied only mapping quality and missing data filters, and included only females in the analyses (Table 7)

Table 7: Estimates based on variant and invariant sites, filtered for mapping quality (40) and missing data (10%), estimated in females only.

| | Estimated in Pixy | | Estimated in ANGSD | |
|---|---|---|---|---|
| | GSL | SSL | GSL | SSL |
| $\pi^{AUT}$ | 0.00028 | 0.00045 | 0.00036 | 0.00082 |
| $\pi^{X}$ | 0.00013 | 0.00023 | 0.00018 | 0.00060 |
| $\pi^{X}$ / $\pi^{AUT}$ | 0.47673 | 0.52463 | 0.50242 | 0.72503 |
| $\boldsymbol{N_e^F/N_e^M}$ | **0.03463** | **0.27071** | **0.15479** | **2.41229** |
| $\theta_w^{\mathrm{AUT}}$ | 0.00043 | 0.00384 | 0.00041 | 0.00125 |
| $\theta_w^{X}$ | 0.00023 | 0.00280 | 0.00027 | 0.00107 |
| $\theta_w^{X}/\theta_w^{\mathrm{AUT}}$ | 0.54415 | 0.72912 | 0.66268 | 0.85316 |
| $\boldsymbol{N_e^F/N_e^M}$ | **0.38335** | **2.49891** | **1.41134** | **9.11529** |

Notably, even small changes in summary statistic between Tables 6 and 7 can have a substantial impact on the $N_e$ ratios.

The ratios derived from ANGSD $\theta_w$ estimates are now consistent with species biology, although GSL ratio still appears underestimated (Table 7). This lower ratio could reflect a true pattern, suggesting that the sex-ratio in GSL may be less biased than previously thought. Such interpretation would be consistent with our other results suggesting female GSL may move more frequently than estimated by behavioural observations or isotope analyses. Alternatively, the lower ratio could be associated with a historical bottleneck experienced by GSL during the colonisation of the archipelago, as statistics are particularly sensitive to changes in effective population size. Similarly, recent decrease in population size have been shown to reduce variation on X (Pool and Nielsen, 2007, Figure S2). This could help explain the consistent underestimation of the $N_e$ ratios based on $\pi$ in GSL whereas those obtained for SSL appear more biologically plausible.

Overall, Pixy estimates are consistently implausible and highly variable, whereas ANGSD estimates are more stable across runs. We therefore kept the ANGSD results from a final run where we additionally applied a minimum depth filter of 2 reads per individual (-setMinDepth2, Table 8)

In summary, our results suggest that $\pi$ may not be the most suitable estimator for inferring $N_e$ ratios as it is highly sensitive to the number of sites and will be strongly affected by data filtering. In our case, recent declines in GSL population size associated with El-Niño events may also reduced diversity on X, leading to underestimation of the $N_e$ ratios. In contrast, $\theta_W$ in our case appears to be more stable and generally presents narrower ranges (Figure 7), although the impact of demographic changes in the deeper past must be taken into account.

Table 8: ANGSD estimates based on variant and invariant sites, filtered for minimum depth of 2, mapping quality and missing data, estimated in females.

| | GSL | SSL |
|---|---|---|
| $\pi^{AUT}$ | 0.00036 | 0.00080 |
| $\pi^{X}$ | 0.00018 | 0.00058 |
| $\pi^{X}$ / $\pi^{AUT}$ | 0.49914 | 0.71862 |
| $\boldsymbol{N_e^F/N_e^M}$ | **0.13865** | **2.28320** |
| $\theta_w^{\mathrm{AUT}}$ | 0.00040 | 0.00114 |
| $\theta_w^{X}$ | 0.00026 | 0.00095 |
| $\theta_w^{X}/\theta_w^{\mathrm{AUT}}$ | 0.65208 | 0.83352 |
| $\boldsymbol{N_e^F/N_e^M}$ | **1.28466** | **7.01638** |

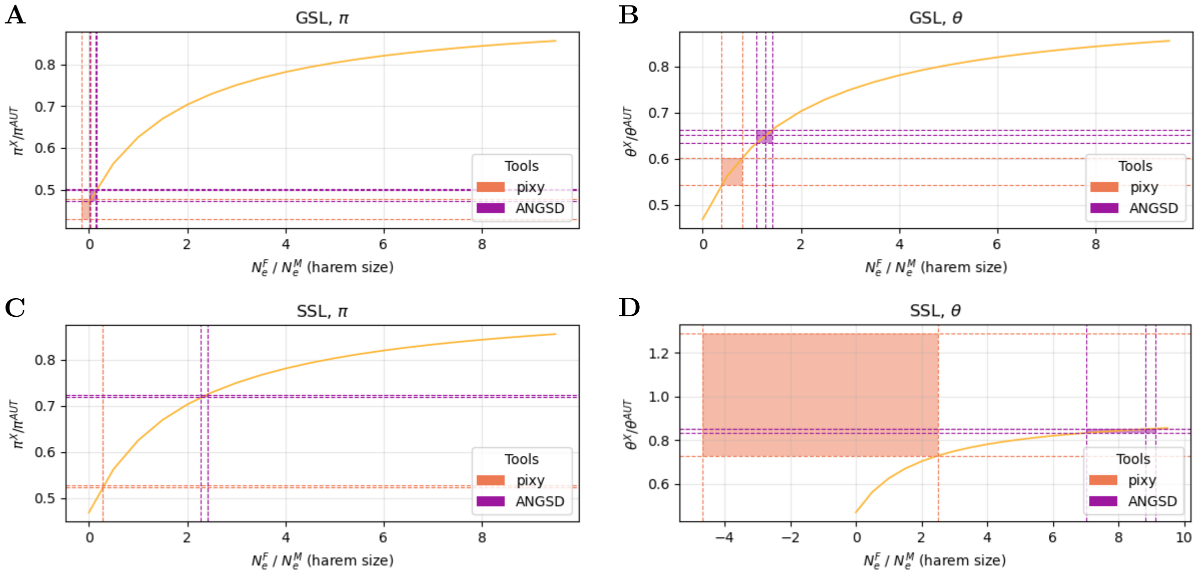

Figure 7: Summary statistics estimates compared to theoretical expectations.

#### 2.4 Effect of downsampling chromosome 9

Although we selected the autosome that was closest in length to the X chromosome, we obtained many more variants in the AUT dataset. This might have led to the difference in signal clarity between the X and AUT, as the population structure in the X does not entirely conform to our expectations ($MT > X > AUT > Y$).

To check whether the noise in population structure could be biased by the different number of SNPs between X and AUT datasets, we used a bash one-liner to randomly downsampled the AUT VCF to the number of X SNPs and repeated the PCA and ADMIXTURE analyses. Those comparisons do not point to number of sites being the source of the lower population structure in X and show similar structure between both baseline and downsampled chromosome 9 datasets (Figures 8 and 9).

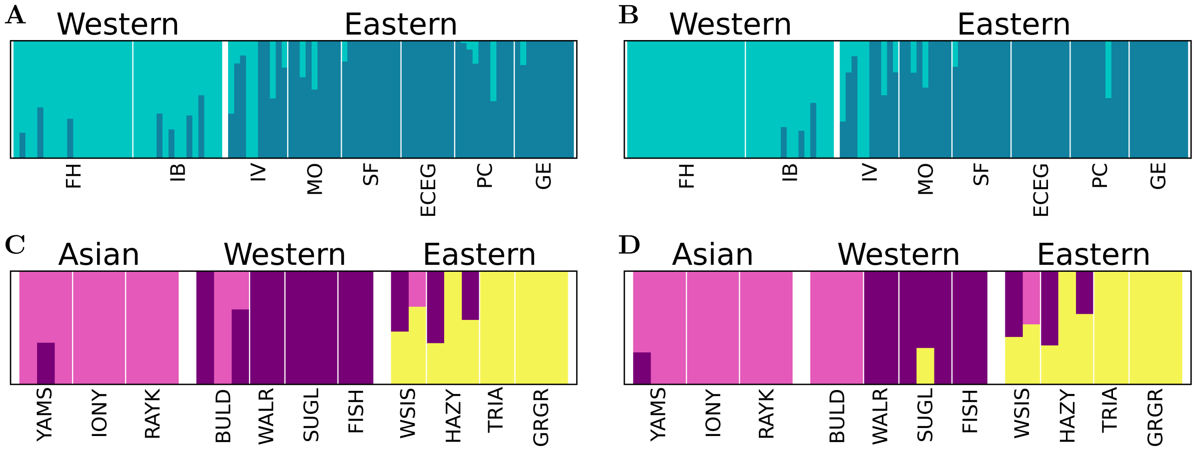

Figure 8: ADMIXTURE proportions comparisons in CM019810.2 for GSL (**A**, **B**) and SSL (**C**, **D**) based on downsampled (**A**, **C**) and baseline data (**B**, **D**).

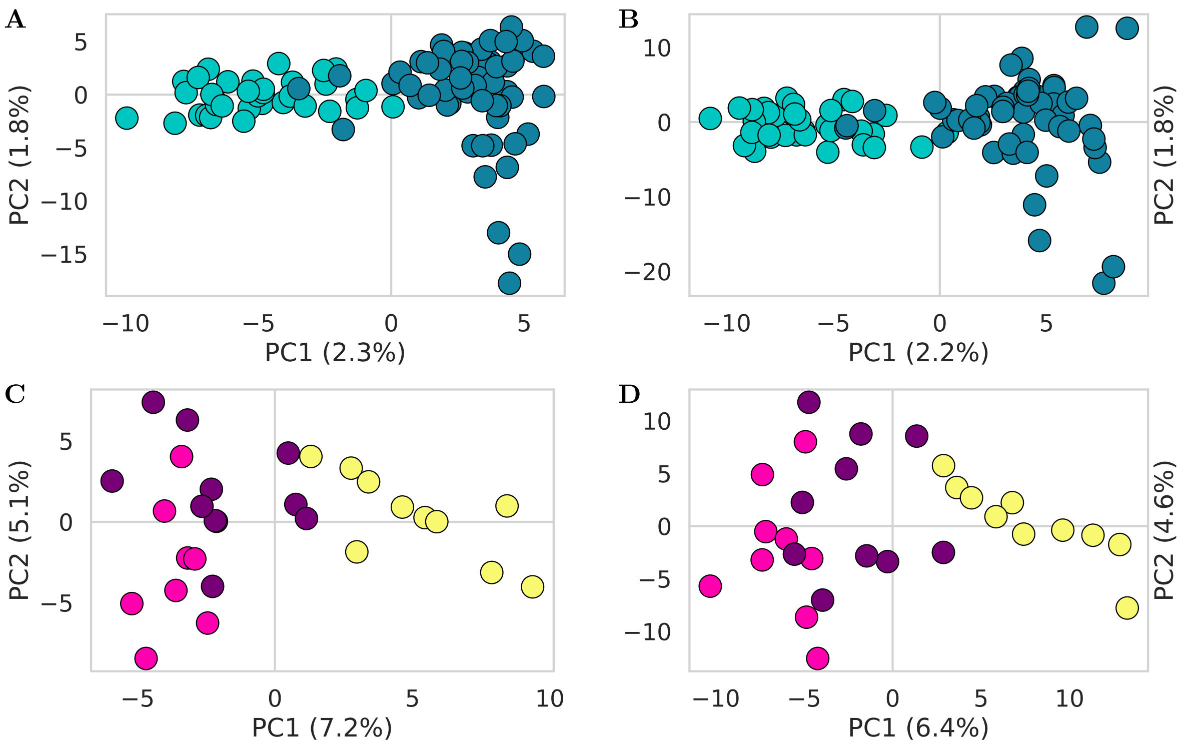

Figure 9: PCA comparisons in CM019810.2 for GSL (**A**, **B**) and SSL (**C**, **D**) based on downsampled (**A**, **C**) and baseline data (**B**, **D**).

#### 2.5 Effect of sequencing coverage on the quality of the variants

Low-coverage sequencing affects variant detection, population structure inference, and estimates of genetic diversity. To evaluate the magnitude of these effects, we performed a coverage sensitivity analysis using 6 female SSL individuals sequenced with coverage of at least 16×. The reads were then randomly downsampled with picard DownsampleSam to generate datasets with an average autosomal coverage of 7×, 10×, and 16× coverage. Each data set was processed using the same variant-calling and filtering pipeline.

**The difference in callable sites and detected variants**

We compared the number of callable positions, SNPs, indels, and singletons recovered from chromosome X (X) and chromosome 9 (AUT) at all three levels of coverage (Table 9).

Table 9: Summary of callable sites and detected variants on chromosome X (X) and chromosome 9 (AUT) under different sequencing coverage thresholds.

| cov | chr | N records | N no-ALTs | N SNPs | N indels | N singletons |
|---|---|---|---|---|---|---|
| 7x | AUT | 47 575 281 | 47 148 106 | 351 137 | 76 038 | 68 120 |
| 10x | AUT | 47 639 842 | 47 225 405 | 337 715 | 76 722 | 49 641 |
| 16x | AUT | 47 713 942 | 47 311 384 | 325 570 | 76 988 | 33 526 |
| 7x | X | 66 174 094 | 65 704 746 | 39 2824 | 76 524 | 90 082 |
| 10x | X | 66 247 418 | 65 802 825 | 36 7583 | 77 010 | 61 468 |
| 16x | X | 66 333 502 | 65 913 432 | 34 2897 | 77 173 | 35 084 |

The number of callable positions, non-variant sites, and indels was mostly insensitive to sequencing coverage, differing by less than 1% between datasets. In contrast, SNP counts showed a greater dependence on coverage, differing by approximately 7%. As expected, singleton variants were the most sensitive category, with counts differing by up to 50%, reflecting the reduced power to detect rare alleles with lower coverage.

**Difference in SNP quality**

In addition, we assessed the effect of coverage on SNPs quality using metrics reported by bcftools: *QUAL* (variant quality score), *DP* (read depth), *RPBZ* (read position bias), *MQBZ* (mapping quality bias), *MQSBZ* (mapping quality versus strand bias), *BQBZ* (base quality bias), *SCBZ* (soft-clipping bias), and *MQ* (mapping quality). The recommended filtering thresholds for these metrics are provided in the samtools/bcftools HTSlib filtering documentation.

Across all coverage levels, the distributions of quality metrics were very similar (Figures 10–12). For all metrics except $QUAL$, the 99% of the data remained within or very close to the recommended filtering thresholds regardless of coverage. The $QUAL$ distribution shifted towards higher values with increasing coverage, reflecting the greater statistical support for SNPs at higher read depths. However, these shifts are unlikely to affect the final set of SNPs, because filtering removes low-quality variants. In general, quality metrics showed only minor differences between coverage levels, suggesting that downsampling primarily affected the power to detect variants rather than the quality of the variants.

**Difference in estimates of genetic diversity**

Finally, we checked the response of the summary statistics and the sex ratio estimates on the different levels of coverage (Table 10). Coverage has a great influence on both summary statistics $\pi$ and $\theta_W$ and, as a consequence, on estimates of the sex ratio, especially for $\theta_W$. Different values of $\alpha$ also have a huge impact, especially when applied to the low-coverage 7x data. The overall tendency between statistics is as expected - the higher the coverage, the closer the $\pi$ and $\theta_W$ estimates become.

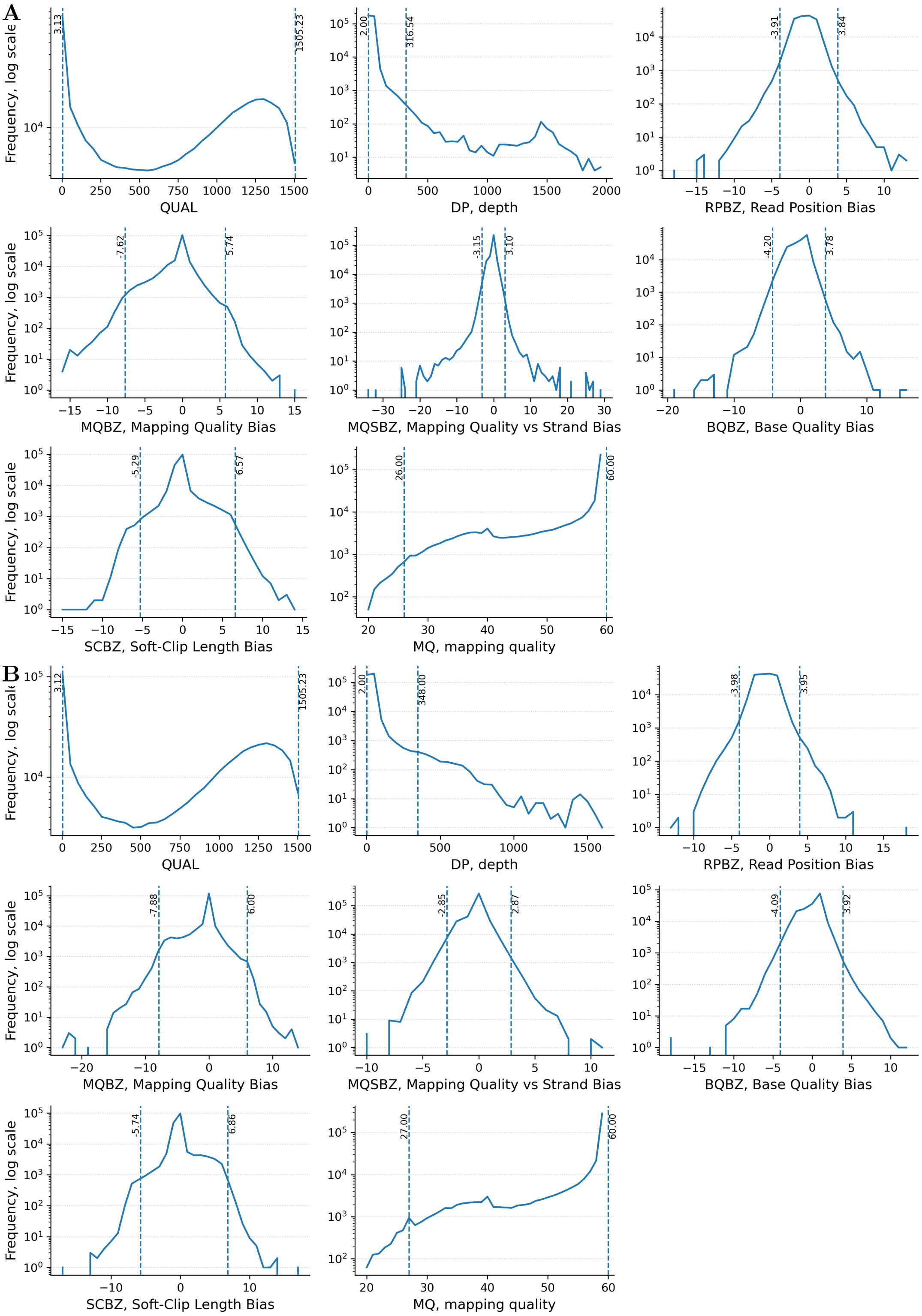

Figure 10: SNPs quality statistics at 7× coverage for the chromosomes 9 (**A**) and X (**B**). Dashed lines show 99% intervals.

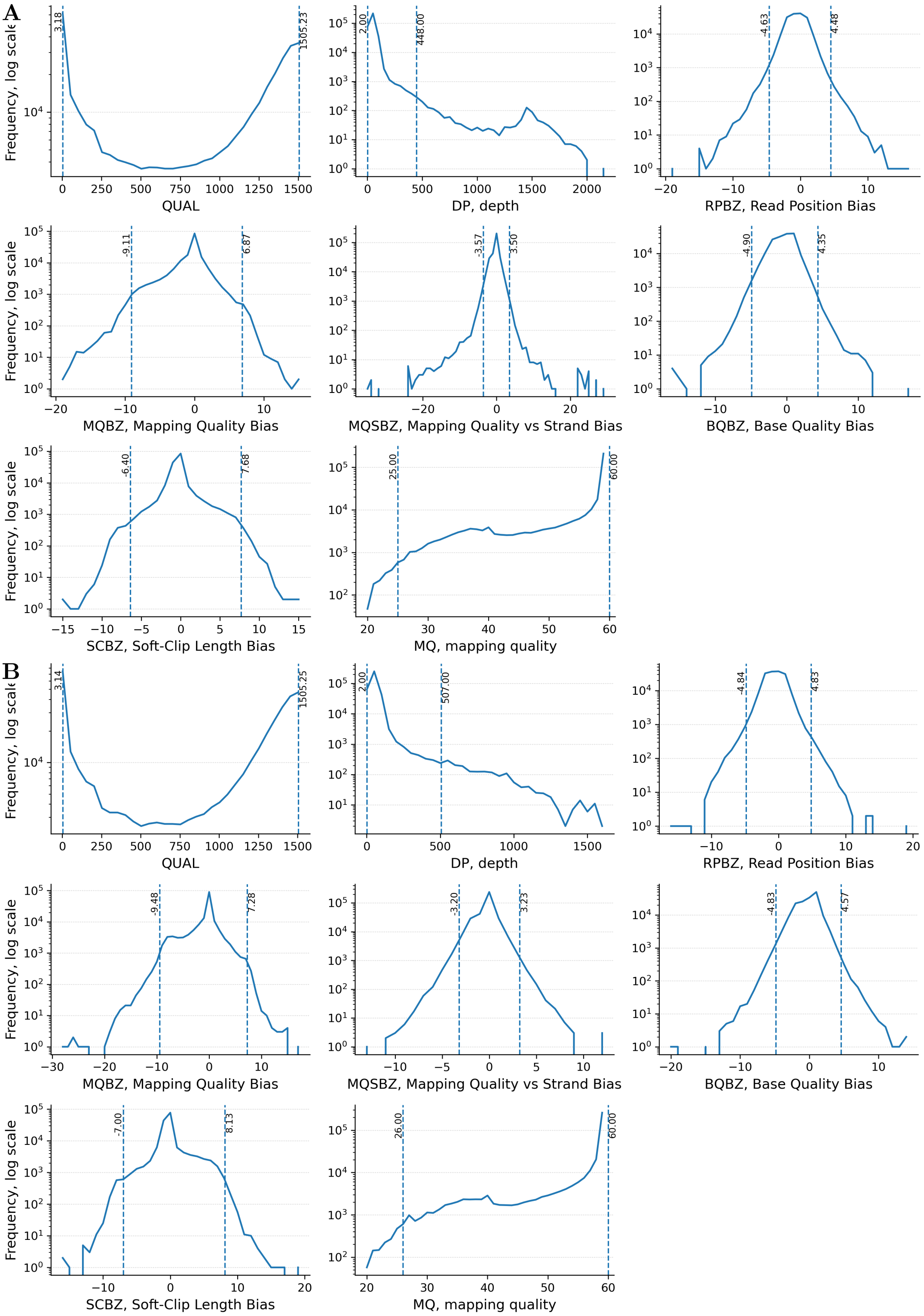

Figure 11: SNPs quality statistics at 10× coverage for the chromosomes 9 (**A**) and X (**B**). Dashed lines show 99% intervals.

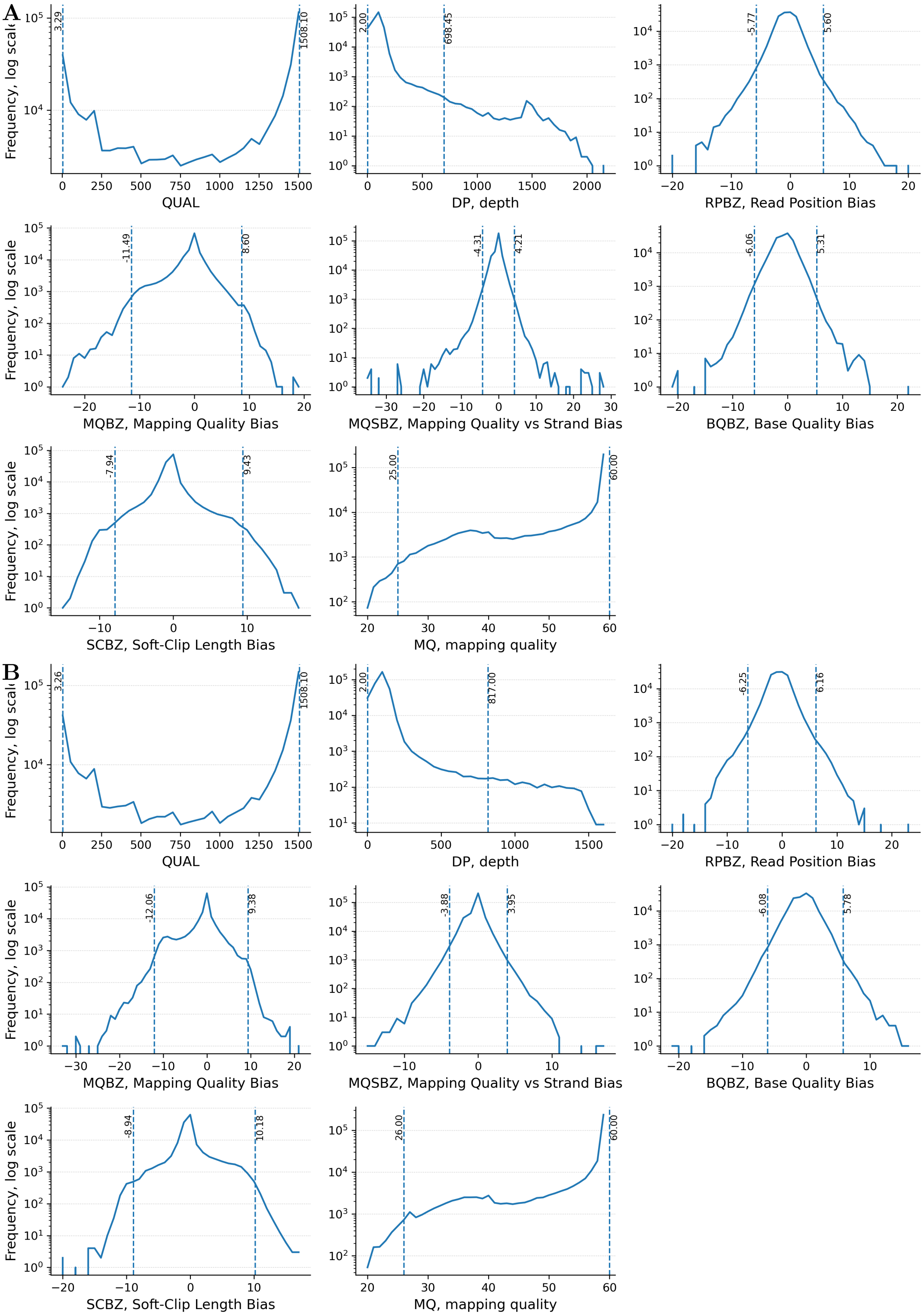

Figure 12: SNPs quality statistics at 16× coverage for the chromosomes 9 (**A**) and X (**B**). Dashed lines show 99% intervals.

Table 10: Summary statistics, calculated with ANGSD ($\pi$ and $\theta_W$) and sex ratio estimates ($N_e^F/N_e^M$), applied to the data of coverage 7x, 10x and 16x, and using $\alpha = 2$ and $\alpha = 3$

| | 7x | 10x | 16x |
|---|---|---|---|
| $\theta_W^{AUT}$ | 0.000881 | 0.0007889 | 0.000544 |
| $\theta_W^X$ | 0.000825 | 0.000711 | 0.000412 |
| $\theta_W^X/\theta^{AUT}$ | 0.936 | 0.902 | 0.758 |
| $N_e^F/N_e^M(\alpha = 2)$ | 13 | 8 | 2 |
| $N_e^F/N_e^M(\alpha = 3)$ | 729 | 24 | 3 |
| $\pi^{AUT}$ | 0.000711 | 0.000664 | 0.000530 |
| $\pi^X$ | 0.000630 | 0.000571 | 0.000408 |
| $\pi^X/\pi^{AUT}$ | 0.885 | 0.861 | 0.769 |
| $N_e^F/N_e^M(\alpha = 2)$ | 6 | 5 | 2 |
| $N_e^F/N_e^M(\alpha = 3)$ | 16 | 10 | 3 |

### 3 Effective population size estimates for males and females.

This section derives expressions for estimating the effective population sizes specific to sex (male $N_e^M$ and female $N_e^F$), combining information from the genetic diversity of AUT and X. The approach relies on well-established relationships between effective population size and the inheritance modes of different genomic compartments.

For AUT, the effective population size reflects contributions from both sexes symmetrically and is given by the following:

$$N_e^{AUT} = \frac{4N_e^F N_e^M}{N_e^F + N_e^M}$$

For X, since it has a sex-biased inheritance pattern (spending $\frac{2}{3}$ of its time in females and $\frac{1}{3}$ in males), the equation is as follows:

$$N_e^X = \frac{9N_e^F N_e^M}{2N_e^F + 4N_e^M}$$

Under the neutral model with constant population size and no migration the ratio of X to AUT effective population size $Q = N_e^X / N_e^{AUT} = 0.75$ with no sex-bias, when female bias (for example, polygyny) $Q > 0.75$, and when male bias (for example, polyandry) $Q < 0.75$

By algebraically rearranging both expressions, $N_e^F$ can be expressed as a function of $N_e^M$ (and vise versa). Equating these alternative expressions allows one to eliminate one variable and solve for the other explicitly. This yields solutions for effective population sizes specific to the sex.

$$N_e^{AUT} = \frac{4N_e^F N_e^M}{N_e^F + N_e^M} \implies N_e^F = \frac{N_e^{AUT} N_e^M}{4N_e^M - N_e^{AUT}}, N_e^M = \frac{N_e^{AUT} N_e^F}{4N_e^F - N_e^{AUT}}$$

$$N_e^X = \frac{9N_e^F N_e^M}{2N_e^F + 4N_e^M} \implies N_e^F = \frac{4N_e^X N_e^M}{9N_e^M - 2N_e^X}, N_e^M = \frac{2N_e^X N_e^F}{9N_e^F - 4N_e^X}$$

$$\frac{N_e^{AUT} N_e^M}{4N_e^M - N_e^{AUT}} = \frac{4N_e^X N_e^M}{9N_e^M - 2N_e^X} \implies \boxed{N_e^M = \frac{2N_e^X N_e^{AUT}}{16N_e^X - 9N_e^{AUT}}}$$

$$\frac{N_e^{AUT} N_e^F}{4N_e^F - N_e^{AUT}} = \frac{2N_e^X N_e^F}{9N_e^F - 4N_e^X} \implies \boxed{N_e^F = \frac{2N_e^X N_e^{AUT}}{9N_e^{AUT} - 8N_e^X}}$$

To make these estimates empirically applicable, the effective population sizes are further expressed in terms of nucleotide diversity ($\theta$) and mutation rates ($\mu$). Under standard population genetic theory, diversity scales with ploidy-specific effective population size and mutation rate:

$$N_e^{AUT} = \frac{\theta^{AUT}}{4\mu^{AUT}}, \quad N_e^X = \frac{\theta^X}{4\mu^X}$$

Substituting these into the expressions above provides formulas for $N_e^M$ and $N_e^F$ directly in terms of observable quantities ($\theta$) and assumed mutation rates ($\mu$).

$$N_e^M = \frac{2\frac{\theta^X}{4\mu^X}\frac{\theta^{AUT}}{4\mu^{AUT}}}{16\frac{\theta^X}{4\mu^X} - 9\frac{\theta^{AUT}}{4\mu^{AUT}}}, \quad N_e^F = \frac{2\frac{\theta^X}{4\mu^X}\frac{\theta^{AUT}}{4\mu^{AUT}}}{9\frac{\theta^{AUT}}{4\mu^{AUT}} - 8\frac{\theta^X}{4\mu^X}}$$

$\mu^{AUT}$ was published by Peart et al., 2020. $\mu^X$ can be derived using the theoretical expectations for $\mu$ as follows:

$$\mu^{AUT} = \frac{\mu^F + \mu^M}{2}, \quad \mu^X = \frac{2\mu^F + \mu^M}{3}$$

By expressing $\mu^M = \alpha\mu^F$ we obtain $\mu^X = \frac{4+2\alpha}{3+3\alpha}\mu^{AUT}$

Combining above we get the final sex ratio expression:

$$\frac{N_e^F}{N_e^M} = \frac{48\theta^X(1+\alpha) - 18\theta^{AUT}(2+\alpha)}{18\theta^{AUT}(2+\alpha) - 24\theta^X(1+\alpha)}$$

Based on published empirical data (Sayres et al., 2011, Amster and Sella, 2016, Foote et al., 2016) on dolphin, killer whale data, human, and great ape which also establish male-mediated gene flow, $\alpha$ is estimated to be in the range of 2 to 3 (Figure S1).

In the model built in Pool and Nielsen, 2007, it's assumed that $\mu^X = \mu^{AUT}$, but in our example we have $\mu^X = \frac{4+2\alpha}{3+3\alpha}\mu^{AUT}$ with $\mu^{AUT} = 5 \times 10^{-9}$ and the estimated $N_e = 12000$ and $N_e = 17000$ and fixed $\alpha = 2$ or $\alpha = 3$ (Figure S2)

### 4 Estimating the sex-biased migration

We estimate the ratio of male to female migration rates ($m^M/m^F$) using genetic differentiation measured in different genomic compartments.

Three following approaches are based on the relationship between genetic differentiation ($F_{ST}$), migration rate ($m$), and effective population size ($N_e$) under a neutral migration-drift equilibrium model.

#### 4.1 Comparing MT and Y differentiation

Following Hedrick et al., 2015, the changes in genetic differentiation between successive generations for females and males can be described as

$$F_{ST,F}^{t+1} = \left[\frac{1}{N_e^F} + \left(1 - \frac{1}{N_e^F}\right) F_{ST,F}^{t}\right](1 - m^F)^2$$

$$F_{ST,M}^{t+1} = \left[\frac{1}{N_e^M} + \left(1 - \frac{1}{N_e^M}\right) F_{ST,M}^{t}\right](1 - m^M)^2$$

Since both compartments have the same mathematical formula, we can first derive a general expression and for simplicity we take $a = \frac{1}{N_e}$. At equilibrium, genetic differentiation remains constant between successive generations:

$$F_{ST}^{t+1} = F_{ST}^{t} = F_{ST}$$

Substituting the above into equations gives:

$$F_{ST} = (a + (1-a)F_{ST})(1-m)^2 = (a(1-F_{ST}) + F_{ST})(1-m)^2$$

$$\frac{F_{ST}}{(1-m)^2} - F_{ST} = a(1 - F_{ST})$$

$$F_{ST}\left(\frac{1}{(1-m)^2} - 1\right) = a(1 - F_{ST})$$

$$\frac{1 - F_{ST}}{F_{ST}} = \frac{\frac{1}{(1-m)^2} - 1}{a}$$

It can be simplified using a Taylor approximation when the migration rate is small.

$$f(x) \approx f(0) + f'(0)x$$

We approximate the function $\frac{1}{(1-m)^2}$.

$$f(m=0) = \frac{1}{(1-0)^2} = 1$$

$$f'(m) = -2\frac{1}{(1-m)^3}(-1) = 2\frac{1}{(1-m)^3}$$

$$f'(m=0) = 2\frac{1}{(1-0)^3} = 2$$

$$\frac{1}{(1-m)^2} \approx 1 + 2m$$

Returning to the equilibrium equation,

$$\frac{1 - F_{ST}}{F_{ST}} = \frac{1 + 2m - 1}{a} = \frac{2m}{a}$$

Substituting $a = 1/N_e$ gives

$$\frac{1 - F_{ST}}{F_{ST}} = 2mN_e \implies m = \frac{1 - F_{ST}}{2F_{ST}N_e}$$

We now apply the general expression to maternally inherited MT and paternally inherited Y.

$$m^{MT} = \frac{1 - F_{ST}^{MT}}{2F_{ST}^{MT}N_e^{MT}}$$

$$m^Y = \frac{1 - F_{ST}^Y}{2F_{ST}^Y N_e^Y}$$

Consequently, for males and females:

$$m^F = \frac{1 - F_{ST}^F}{2F_{ST}^F N_e^F}$$

$$m^M = \frac{1 - F_{ST}^M}{2F_{ST}^M N_e^M}$$

And finally the ratio of male to female migration rates:

$$\frac{m^M}{m^F} = \frac{(1 - F_{ST}^M)}{F_{ST}^M} \frac{F_{ST}^F}{(1 - F_{ST}^F)} \frac{N_e^F}{N_e^M}$$

#### 4.2 Comparing MT and AUT differentiation

An alternative approach is to compare MT and AUT differentiation, following Hedrick et al., 2015 The equation for AUT is:

$$F_{ST}^{t+1} = \left[\frac{1}{2N_e} + \left(1 - \frac{1}{2N_e}\right) F_{ST}^t\right] (1 - m)^2$$

Unlike MT and Y, AUT are inherited biparentally. Consequently, AUT differentiation is influenced by migration and genetic drift in both sexes. We can therefore repeat the derivation from the previous section, this time using genetic drift for diploid populations: $a = \frac{1}{2N_e}$

At migration-drift equilibrium, we obtain:

$$\frac{1 - F_{ST}^{AUT}}{F_{ST}^{AUT}} = 4m^{AUT} N_e^{AUT}$$

Autosomes are inherited equally from the mother and father, thus the AUT migration rate is the average of the female and male migration rates:

$$m^{AUT} = \frac{m^M + m^F}{2}$$

Additionally, under the standard two-sex population model, the AUT effective population size is:

$$N_e^{AUT} = \frac{4N_e^M N_e^F}{N_e^M + N_e^F}$$

We now combine the equilibrium equations for AUT and MT differentiation:

$$\frac{(1 - F_{ST}^{AUT})}{F_{ST}^{AUT}} \frac{F_{ST}^{MT}}{(1 - F_{ST}^{MT})} = 4\frac{4N_e^M N_e^F}{N_e^M + N_e^F} \frac{m^M + m^F}{2} \frac{1}{2m^F N_e^F}$$

$$\frac{(1 - F_{ST}^{AUT})}{F_{ST}^{AUT}} \frac{F_{ST}^{MT}}{(1 - F_{ST}^{MT})} = 4\frac{N_e^M}{N_e^M + N_e^F} \left(1 + \frac{m^M}{m^F}\right)$$

And finally get the ratio of male to female migration rates:

$$\frac{m^M}{m^F} = \frac{1}{4}\frac{(1 - F_{ST}^{AUT})}{F_{ST}^{AUT}} \frac{F_{ST}^{MT}}{(1 - F_{ST}^{MT})} \left(1 + \frac{N_e^F}{N_e^M}\right) - 1$$

#### 4.3 Comparing X and AUT differentiation

Following Ramachandran et al., 2008, we used the relative differentiation of X and AUT to estimate male-biased migration. Under the two sex infinite-island model, the effective numbers of female and male migrant per deme per generation are:

$$M^F = \frac{1}{1 - r}\left[\frac{2 - r}{6F_{ST}^X} - \frac{1}{8}\left(\frac{1}{F_{ST}^{AUT}} + \frac{1}{3}\right) - \frac{1 - r}{6}\right]$$

$$M^M = \frac{1}{r}\left[-\frac{2-r}{6F_{ST}^X} + \frac{1}{4}\left(\frac{1}{F_{ST}^{AUT}} + \frac{1}{3}\right) - \frac{r}{6}\right],$$

where $r = \frac{N_e^F}{N_e^F + N_e^M}$ is the female fraction of the population size, and $M^F = N_e^F m^F$ and $M^M = N_e^M m^M$.

We approximated $r$ from the estimated female-to-male effective population size ration as:

$$r = \frac{N_e^F/N_e^M}{1 + N_e^F/N_e^M}$$

Using the observed $F_{ST}^{AUT}$, $F_{ST}^X$ between 2 stocks, and $r$, we obtained a male-to-female migration ratio $M^M/M^F = 6.19$, consistent with the previously obtained results based on $AUT$ and $MT$. For SSL, however, the value was negative due to $M^F < 0$, resulting in $M^M/M^F = -7.079$. The negative value of migrants is impossible, thus the obtained value cannot be interpreted as either the direction or magnitude of migration. Instead it indicates that the observed combination of $F_{ST}^{AUT}$, $F_{ST}^X$, and $r$ lies outside of the parameter space of the model. In addition, all the estimates based on pairwise $F_{ST}$ values among the three stocks were also negative.

As stated in the model description, for $M^F$ to be positive, the $F_{ST}^X$ must satisfy:

$$\frac{2F_{ST}^{AUT}(2-r)}{3 + F_{ST}^{AUT}(1-2r)} < F_{ST}^X < \frac{4F_{ST}^{AUT}(2-r)}{3 + F_{ST}^{AUT}(5-4r)}$$

Given the observed SSL values of $F_{ST}^{AUT}$ and $r$, $F_{ST}^X$ should be $0.016 < F_{ST}^X < 0.0318$, with the observed value $F_{ST}^X = 0.03245$, which is only slightly above the boundary.

Thus, only slight difference in parameters separates the results from interpretability. The model is highly sensitive to small changes in input parameters. Uncertainty may arise starting from sequencing errors to variant filtering. Moreover the model assumes neutral migration-drift equilibrium, infinite-island model and same female fraction in each deme. As discussed in the main text, these assumptions are unlikely to be fully met in our study model.

Overall, the current model provides an additional confirmation of male biased migration for GSL, whereas the SSL data yield an incompatible parameter combination and do not provide an interpretable result.
